# Enhancing and stabilizing a high-yield industrial *Rhodobacter sphaeroides* strain for coenzyme Q_10_ production

**DOI:** 10.1101/2025.07.16.665108

**Authors:** Mindong Liang, Xinwei He, Dongyuan lv, Jing Liu, Kefeng Wang, Yingrui Hou, Weishan Wang, Linquan Bai, Guang Liu, Zhichun Zhu, Dan Li, Biqin Chen, Lixin Zhang, Gao-Yi Tan

## Abstract

Strengthening high-yield phenotypes while maintaining physiological and genetic stability presents a significant challenge in the improvement of high-yield industrial strains (HIS). Coenzyme Q_10_ (CoQ_10_), a crucial quinone electron carrier in the electron transport chain, is widely used in the prevention and treatment of cardiovascular diseases. In this study, the established HIS *Rhodobacter sphaeroides* HY01, employed for CoQ_10_ production, was engineered to enhance productivity while ensuring strain stability. Comparative omics identified the PrrAB two-component system as an oxygen-responsive regulator that links CoQ_10_ biosynthesis to photosynthetic pathways. Mutagenesis of PrrA, guided by AlphaFold3 modeling and fluorescence screening, introduced mutations that led to a 37.5% increase in CoQ_10_ production. To address phenotypic reversion due to metabolic burden, genome-scale CRISPR interference (CRISPRi) screening identified key genes involved in DNA repair and stress adaptation. Deletions of these genes generated a stable strain that achieved 3.6 g/L CoQ_10_ in a 50- L pilot-scale fed-batch fermentation, surpassing previous reports. This study reveals PrrAB-mediated flux partitioning for redox homeostasis and provides a framework for stabilizing burdened phenotypes in photosynthetic microbes, advancing the sustainable production of redox-active metabolites.

**Bullet points:**

1. Identified the PrrAB two-component system as a critical global regulator of CoQ_10_ biosynthesis in *Rhodobacter sphaeroides*.
2. PrrA was evolved through fluorescence-based screening and rational protein engineering, significantly enhancing CoQ_10_ biosynthesis in industrial high-yield strain.
3. Genome-scale CRISPRi screening identified genes affecting *R. sphaeroides* HY01 stability enabling targeted modifications to stabilize high-yield CoQ_10_ phenotype.
4. Achieved record 3.6 g/L CoQ_10_ yield in 50-L pilot-scale bioreactors enhancing microbial productivity stabilizing high-yield phenotypes advancing strain engineering.

## Introduction

High-yield industrial strains (HISs) undergo extensive optimization through diverse breeding and engineering strategies to produce valuable compounds, including pharmaceuticals like avermectin ^1^, fine chemicals such as porphyrins ^2^, and nutraceuticals like coenzyme Q_10_ (CoQ_10_) ^3^. Yet, the high-yield phenotype imposes substantial metabolic burdens or stress ^4-6^, limiting the transferability of strategies effective in wild-type or low-yield strains. Genetic and/or physiological instability further complicates matters, as interventions such as gene knockouts or overexpression can enhance yields but often trigger phenotypic reversion, undermining long-term stability ^7-9^. Balancing productivity with strain stability thus remains a core challenge in strain improvement or breeding for industrial microbiology.

CoQ_10_ serves as a vital quinone electron carrier in bacterial and mitochondrial electron transport chains, where its deficiency disrupts redox homeostasis, contributing to mitochondrial disorders, neurodegeneration, and cardiovascular diseases ^10^. This has driven its widespread use as a supplement for mitigating oxidative stress and preventing cardiovascular diseases ^11^. The global CoQ_10_ market was valued at USD 614.2 million in 2022 and is projected to grow at a CAGR (Compound Annual Growth Rate) of 10.2%, reaching USD 1.33 billion by 2030, according to Grand View Research (www.grandviewresearch.com). China plays a leading role in the global supply of CoQ_10_ raw materials, but tightening environmental regulations necessitate HIS engineering to improve efficiency and reduce ecological footprints ^12^.

Microorganisms like *Agrobacterium tumefaciens* and *Rhodospirillum rubrum* excel in CoQ10 biosynthesis ^13, 14^, with the photosynthetic bacterium *Rhodobacter sphaeroides* emerging as a premier industrial chassis due to its flexible metabolic modes, including photosynthetic and respiratory pathways ^15, 16^. *R. sphaeroides*’ ability to switch between photosynthetic and respiratory modes under varying oxygen levels makes it ideal for CoQ_10_ production ^17^, yet exposes it to redox imbalances that exacerbate instability ^18^. Prior efforts have targeted physiological roles and structure of CoQ_10_, focusing on redox regulation, isoprenoid precursor enhancement, and pathway optimization ^19-21^. Strategies including MEP (methylerythritol phosphate) pathway overexpression, carotenoid competition blockade, and p-hydroxybenzoate supply augmentation have boosted CoQ_10_ yields in *R. sphaeroides*^22-24^. In our prior work with established HIS *R. sphaeroides* HY01, conventional strategies underperformed ^3^, prompting synthetic promoter engineering for key gene expression ^25^, which elevated CoQ_10_ but revealed instability via spontaneous mutations during prolonged fermentation ^18, 26^. While random mutagenesis has yielded gains ^27, 28^, integrated approaches for yield and stability in HISs remain scarce.

Here, we dissected regulatory intricacies, productivity, and stability in CoQ_10_-producing HISs. Omics analyses revealed key regulators of CoQ_10_ biosynthesis, including the PrrAB two-component system linking it to photosynthetic physiology. Applying global transcription machinery engineering (gTME) ^29^ and saturation mutagenesis, we used a fluorescence reporter to enable high-throughput screening of PrrA variants for CoQ_10_ yield gains, while genome-scale CRISPRi with phenotypic selection identified genes conferring stress resilience. This yielded a stable strain with unprecedented CoQ_10_ titers in pilot-scale bioreactors, advancing insights into microbial redox homeostasis and adaptation mechanisms for sustainable biomanufacturing.

## Results

### Discovery of the critical role of the PrrAB two-component system in CoQ_10_ biosynthesis

In prior plate cultivation of the HY01 strain, we isolated a spontaneous mutant, HY02, displaying a white colony phenotype rather than HY01’s green (Fig. 1a). Fermentation assays showed HY02 produced 82% less CoQ_10_ than HY01, even below wild-type *R. sphaeroides* 2.4.1 levels ^22^. Comparative transcriptomics during shake-flask fermentation uncovered extensive changes in HY02: 1,208 genes downregulated and 251 upregulated at 24 h, shifting to 358 downregulated and 786 upregulated at 48 h (Fig. 1b). Gene Ontology enrichment highlighted involvement in electron carrier activity, respiratory chain components (*e.g.*, NADH dehydrogenase, cytochrome c oxidase), photosynthetic pigment biosynthesis (bacteriochlorophyll and terpenoid pathways), and nitrogen fixation, among others (Fig. 1c; Fig. S1).

**Fig. 1.**
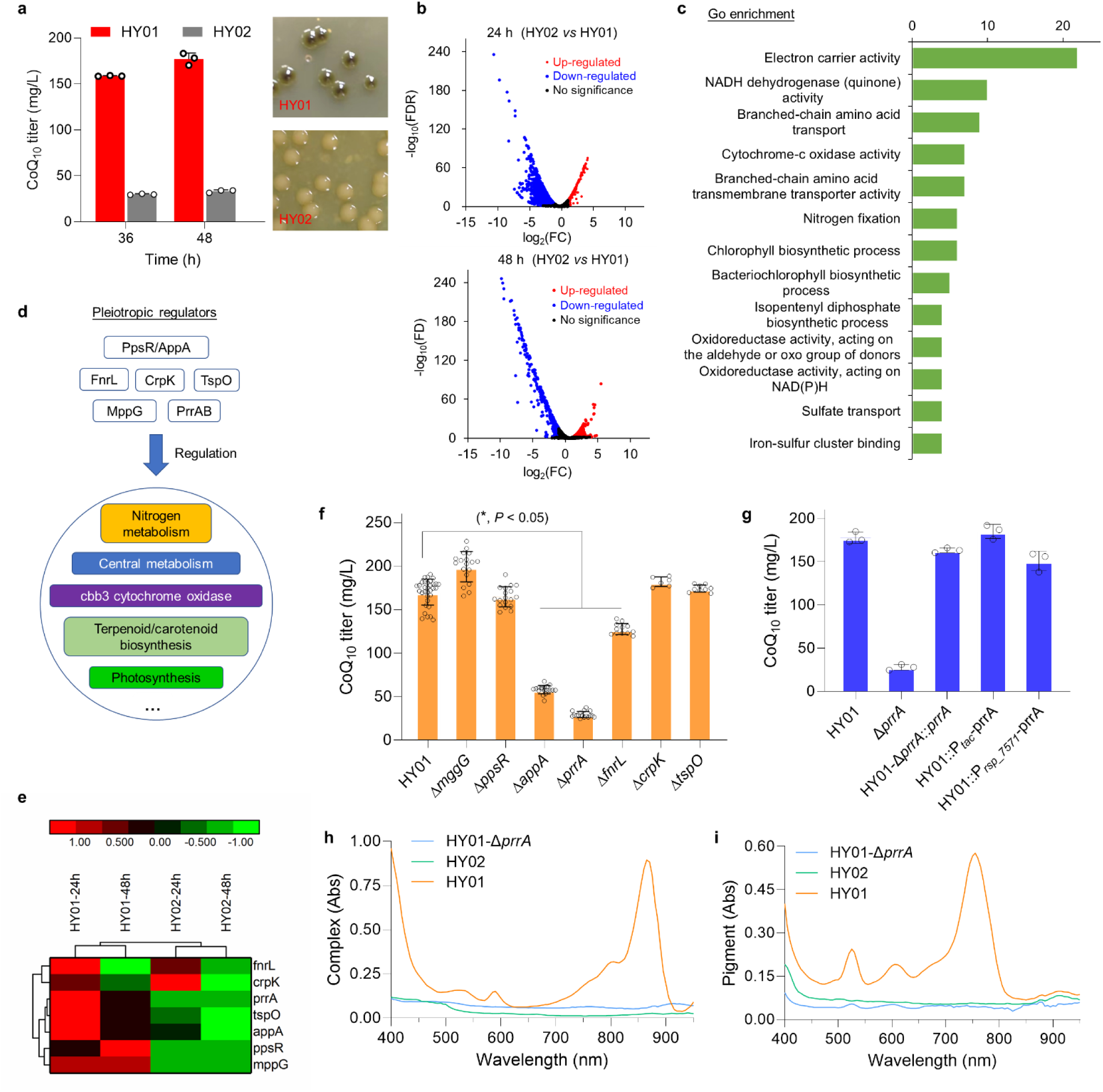
Identification of a key regulatory gene for CoQ_10_ biosynthesis via comparative transcriptomic and genetic analyses. **a,** Colony phenotypes and Q10 production in HY01 and HY02. **b,** Transcriptomic differences between HY01 and HY02 (Volcano Plot; 24h and 48h). **c,** Gene Ontology (GO) analysis of differentially expressed genes. **d,** Identified global regulatory genes in *R. sphaeroides*. **e,** Transcription of regulatory genes in HY01 and HY02. **f,** Effects of different regulatory gens on CoQ_10_ production. **g.** Effects of *prrA* complementation with promoters varying strengths on CoQ_10_ production in *prrA*- inactivated mutant. **h,** Spectral analysis of light-harvesting protein complexes in *prrA*-inactivated HY01. **i.** Spectral analysis of photosynthetic pigments in *prrA*-inactivated HY01. Bar graph with error bars represents mean ± s.d. \**P* < 0.05.

Prior work in *R. sphaeroides* 2.4.1 identified regulators like PpsR/AppA, FnrL, CrpA, TspO, MppG, and PrrAB in photosynthesis, nitrogen metabolism, respiration, and terpenoid synthesis ^30, 31^. Transcriptomics showed downregulation of these in HY02 (Fig. 1d-e). Systematic inactivation of these genes in HY01 reduced CoQ_10_ significantly (*P* < 0.05) only for *fnrL*, *appA*, and *prrA* deletions, with *prrA* knockout (HY01-Δ*prrA*) causing an 84% drop and white phenotype mimicking HY02 (Fig. 1f). Promoter-tuned *prrA* complementation in HY01-Δ*prrA* restored yields (Fig. 1g), while *prrA* duplication in HY02 raised CoQ_10_ to 86 mg/L (Fig. S2), though not fully to HY01 levels, implying additional modulators.

These data identify PrrAB as a key regulator of CoQ_10_ production. Given the white colonies of HY02, we examined the photosynthetic apparatus. The absorbance spectra for HY02 and HY01-Δ*prrA* (Fig. 1h-i) showed a lack of peaks corresponding to the reaction center-light harvesting antenna 1 (RC-LH1) core complexes ^32^, bacteriochlorophyll ^33^, and associated pigments, which coincided with the decline in CoQ_10_ production, hinting at a possible connection between these pathways.

### Red fluorescence-based genetic circuit identifies functional residues in PrrA

PrrAB regulates photosynthesis genes oxygen-dependently in *R. sphaeroides*; *prrA* loss abolishes complexes ^34^. Transcriptomics confirmed bacteriochlorophyll gene downregulation in HY01-Δ*prrA* (Fig. S3). This study examined various HY01-derived strains with differing CoQ_10_ yields cultured in shake flasks. CoQ_10_ and bacteriochlorophyll levels were measured during fermentation, revealing a positive correlation between the two (n = 135, R^2^ = 0.78; Fig. 2a). Bacteriochlorophyll contributed to the green phenotype of the colonies, suggesting that it could be explored as a reliable proxy for CoQ_10_ productivity.

**Figure 2.**
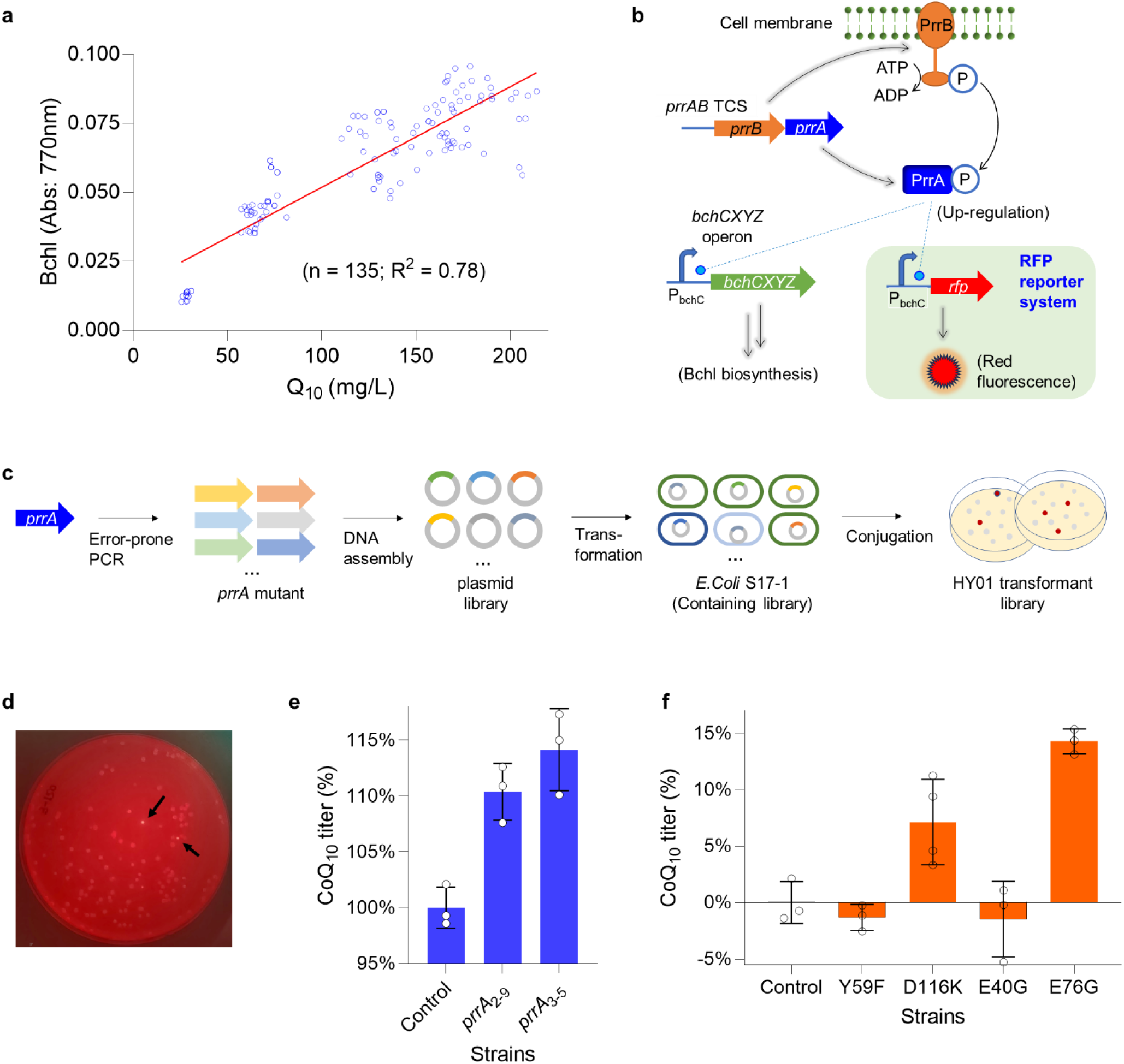
Engineering and screening of *prrA* mutants using a fluorescent reporter system to enhance coenzyme q10 production in *R. sphaeroides*. **a.** Correlation analysis between bacteriochlorophyll content and CoQ10 production in *R. sphaeroides*. **b.** Design of a red fluorescent reporter system driven by promoters of key bacteriochlorophyll biosynthesis genes regulated by the PrrAB two-component system. **c.** Workflow for screening strains combining random point mutagenesis of *prrA* and the fluorescence reporter system. **d.** Isolation of single colonies harboring desired *prrA* mutations using the fluorescence reporter system. Arrows indicate colonies with enhanced red fluorescence. **e.** Effects of selected *prrA* mutants on CoQ10 production. **f.** Contributions of different *prrA* point mutations to improved CoQ10 production. Bar graph with error bars represents mean ± s.d. \**P* < 0.05.

Leveraging the correlation between bacteriochlorophyll and CoQ_10_ accumulation, we engineered an RFP reporter system (Fig. 2b) driven by PrrAB-regulated *bchC* promoters for high-throughput strain screening. The *bchCXYZ* operon is a critical metabolic node regulating bacteriochlorophyll biosynthesis ^35^, and the transcriptional activity of the *bchC* promoter directly affects bacteriochlorophyll production. This system correlates fluorescence changes induced by PrrA mutations with *bchCXYZ* transcription, enabling the identification of clones with enhanced bacteriochlorophyll biosynthesis. The fluorescence-based proxy for bacteriochlorophyll upregulation was designed to enable efficient screening of strains for potential improvements in CoQ_10_ yields.

The workflow (Fig. 2c) involved error-prone PCR library cloning into an integrative vector, conjugation into HY01- Δ*prrA*, and selection of fluorescent colonies with markedly enhanced fluorescence (Fig. 2d), followed by cultivation in 48-well deep-well plates to assess CoQ_10_ contents. Using this method, *prrA*_2-9_ and *prrA*_3-11_ mutants enhancing CoQ_10_ yields were identified (Fig. 2e). Sequencing uncovered Y59F, D116K, E100G, and E76G mutations (Fig. 2f), and single-site validation confirmed D116K and E76G as key mutations, with E76G yielding approximately 15% higher yields compared to HY01. Thus, the RFP-coupled gTME approach targets PrrA to enhance CoQ_10_ production, although further improvements are needed for industrial-scale applications.

### Computational modeling and PrrA functional residues saturation mutagenesis

AlphaFold3 modeling of D116K and E76G variants (Fig. 3a-b; Fig. S4) showed PrrB dimer interfacing two PrrA monomers. E76 forms a salt bridge with PrrB R314 (3.054 Å distance); glycine substitution disrupts it, lowering inter-protein TM-score (ipTM) from 0.44 to 0.41. D116K introduces a hydrogen bond with PrrB R255 (distance 3.439 Å from 4.997 Å), yielding ipTM 0.43. These alterations likely tune phosphotransfer by altering interface dynamics.

**Figure 3.**
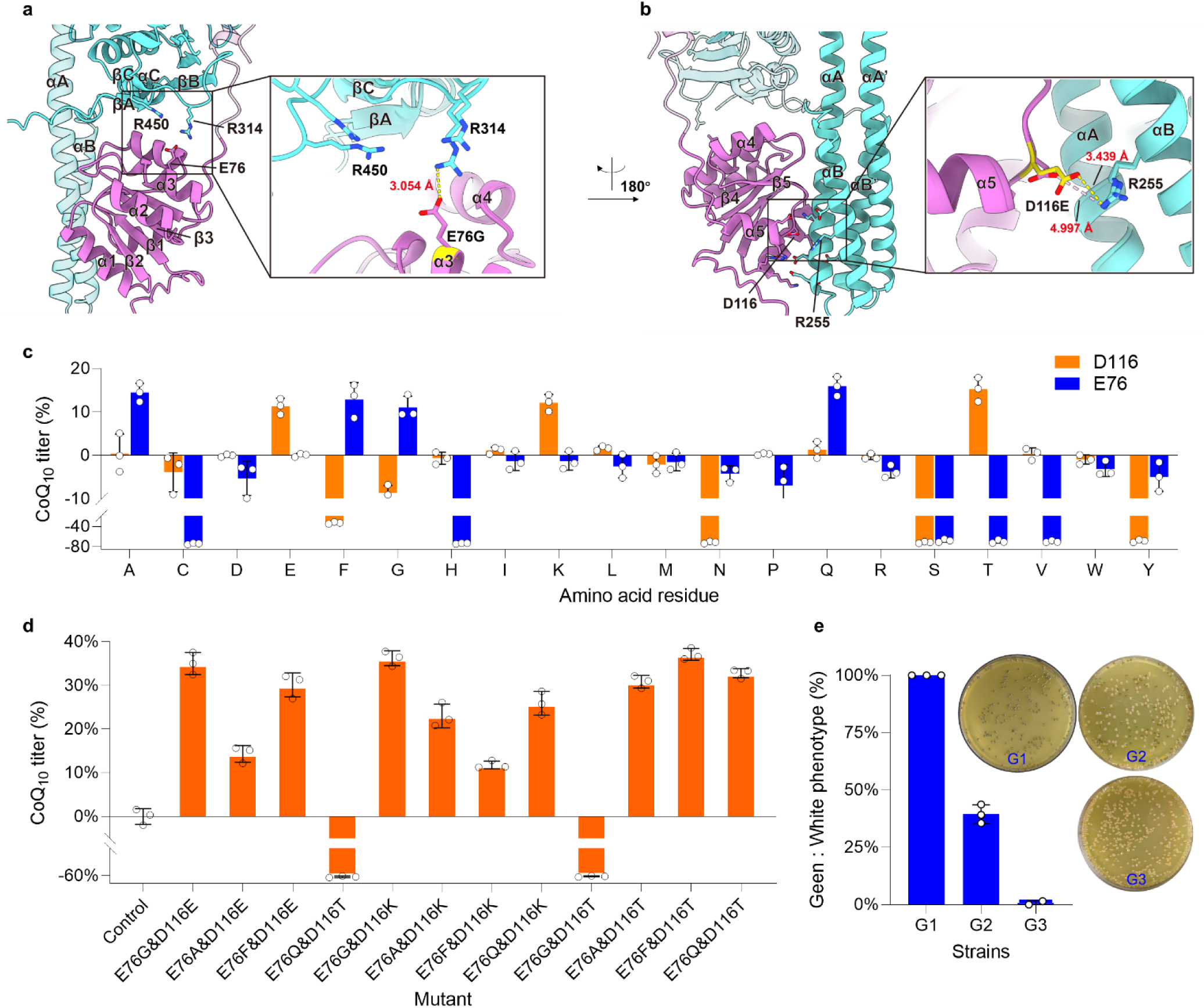
Structural modeling and mutational analysis of PrrA variants affecting CoQ_10_ production and strain stability. **a-b,** AlphaFold3-based structural models of PrrA variants illustrate the potential impact of E76 and D116 mutations on the PrrAB two-component signaling interface. **c,** Effects of saturation mutagenesis at residues E76 and D116 on CoQ_10_ production. **d,** Effects of combinatorial mutations at E76 and D116 on CoQ_10_ yield. **e,** Effects of serial passaging on the phenotypic stability of the engineered strains.

Saturation mutagenesis of the E76 and D116 sites (Fig. 3c) identified variants like E76A/F/G/Q and D116E/K/T, which led to over 10% improvements. Combinations (*e.g.*, E76G&D116E/K, E76F&D116T) synergistically enhanced yields by more than 30% in 48-well plates. The E76F&D116T variant (HY11) achieved a 37.5% increase over HY01 (243 mg/L; Fig. 3d), but when tested for fermentation performance in shake-flask experiments, it showed only a 21.4% increase (Fig. 5a), highlighting scalability issues.

Further analysis revealed that liquid fermentation of HY11 spontaneously produced white variants, with a loss of pigmentation and no further accumulation of CoQ_10_ upon serial passaging. By the third round, full reversion occurred (Fig. 3e; Fig. S5), and colonies with a green phenotype were not observed on agar. This demonstrates engineering of PrrA enhances CoQ_10_ but compromises stability under metabolic stress, necessitating coupled robustness strategies.

### CRISPRi screening for genes enabling stable high-yield industrial strains

Since the accumulation of CoQ_10_ is accompanied by the green colony phenotype in HY01, we designed the strategy shown in Fig. 4a to identify potential targets that could stabilize the high-yield phenotype. Genome-scale CRISPRi screening used a crRNA library in modified pBBR-dCas12a ^2^, conjugated into HY11 (500-1,000 conjugants/experiment). Conjugants were pooled, subcultured in TSB medium (10% inoculation), induced with IPTG, and transferred to CoQ_10_ medium for 72 hours of fermentation. The resulting cultures were diluted and plated to collect green colonies. From ∼20,000 colonies across 20 rounds, 41 stable candidates emerged. crRNA sequencing identified 10 targets (Fig. 4b), with RSP_1864 and RSP_1085 enriched (20 and 9 hits). Annotations implicated DNA replication/repair (e.g., RSP_1028 DNA polymerase I, RSP_0524 Ku protein, RSP_2679 LigD polymerase module) and five hypothetical proteins, including WGR-domain RSP_1085 (Fig. 4c).

**Figure 4.**
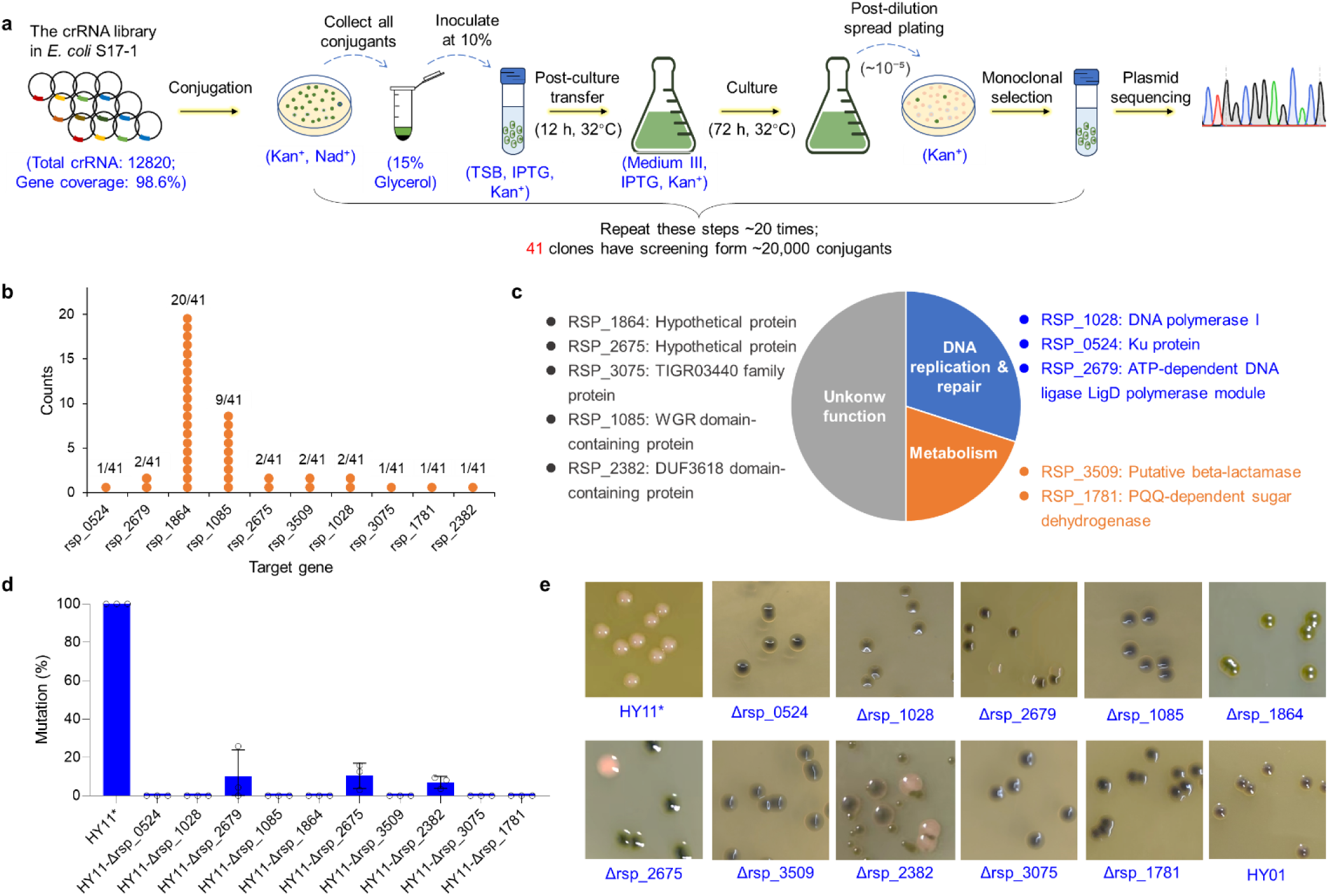
Identification and validation of genes for green colony phenotype using genome-scale CRISPRi screening. **a,** Workflow for identifying genes involved in maintaining green colony phenotype using a genome-wide crRNA library. **b-c,** Candidate genes and their predicted biological functions identified from 41 green phenotype isolates. **d,** Mutation frequency after three rounds of liquid fermentation in HY11 strains with single knockouts of 10 candidate genes. **e,** Colony phenotype and pigmentation of HY11 strains with individual disruptions of identified genes.

To assess the roles of these genes in phenotypic stability, we individually deleted each of the ten candidates in the HY11 strain and evaluated the mutants for spontaneous phenotype switching, marked by white colony emergence, after three serial liquid fermentations (Fig. S5a). Deletions of RSP_2679, RSP_2675, and RSP_2382 yielded white colony frequencies <10%, whereas the other seven mutants produced none (Fig. 4d), indicating that these deletions promote green phenotype maintenance.

Colony morphology analysis revealed that all mutants retained green pigmentation to varying extents but differed in size and hue (Fig. 4e). For example, ΔRSP_1864 and ΔRSP_2675 mutants exhibited yellow-green and grass-green coloration, respectively; deletions of RSP_0524, RSP_1085, RSP_3075, and RSP_1864 produced darker pigmentation than HY11; and ΔRSP_2382 mutants formed mixed large and small colonies. These results suggest complex mechanisms governing gene effects on colony color, yet their inactivation enhances overall phenotypic stability.

### High-yield phenotype stabilization and pilot-scale fermentation of CoQ10 in a 50L bioreactor

To develop a stable, high-yield strain for industrial fermentation, we assessed ten mutants in shake flasks. Some deletions decreased titers by 6-15% vs. HY11 (Fig. 5a), whereas ΔRSP_1028, ΔRSP_1085, ΔRSP_3509, and ΔRSP_3075 maintained or slightly raised them. Double deletions (RSP_1028&RSP_1085, RSP_1028&RSP_3509, RSP_1085&RSP_3509) elevated production by 13%, 7%, and 11% over HY11 (37.2%, 30%, 34.5% vs. HY01; Fig. 5b), showing combinations stabilize and boost the phenotype.

**Figure 5.**
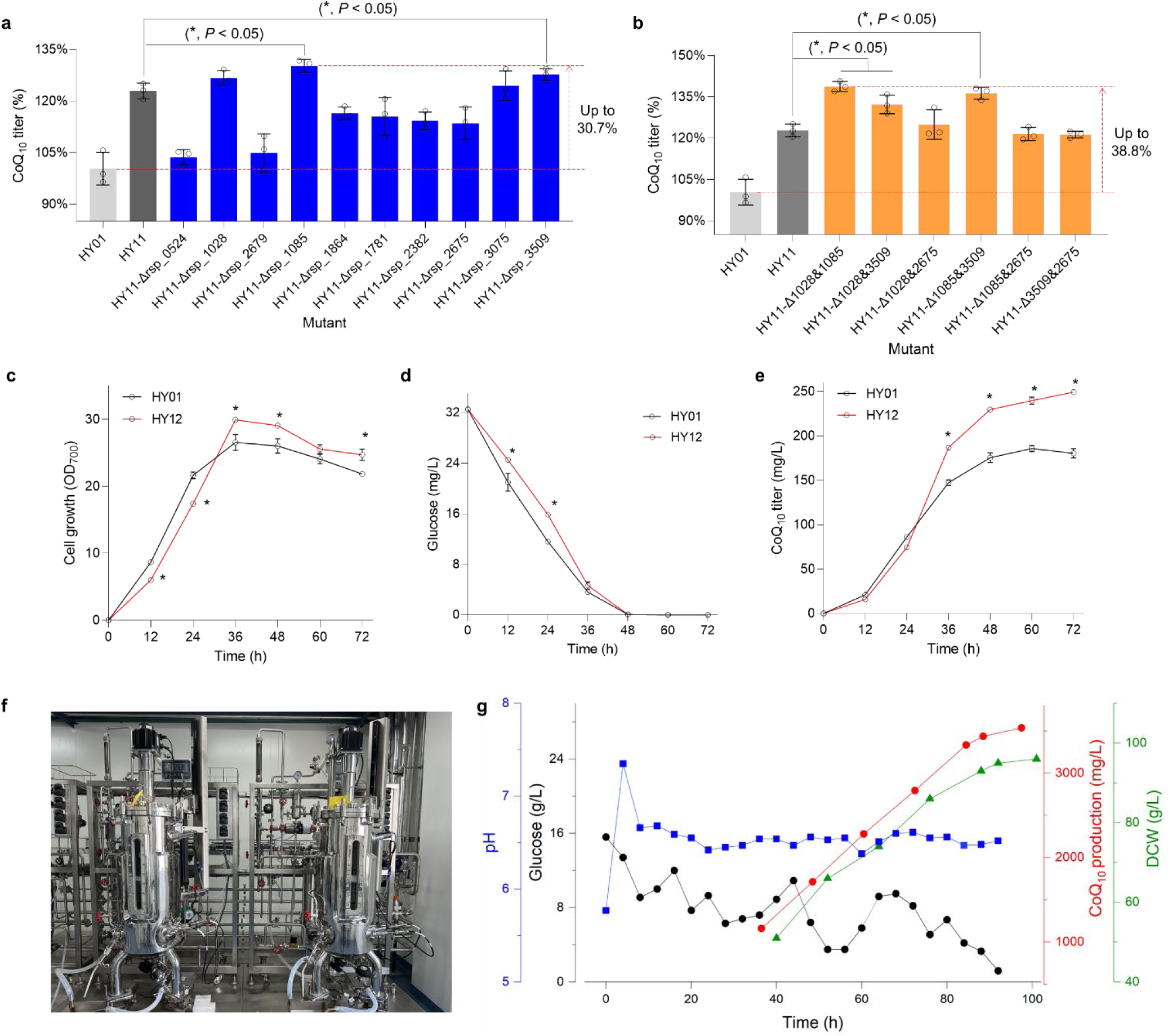
Combinatorial gene deletions enhance CoQ_10_ production and fermentation performance of HY12. **a,** Effects of individual knockout of ten candidate genes on CoQ_10_ production in strain HY11. **b,** Improvement of CoQ_10_ production through combinatorial double-gene deletions. **c–e,** Time-course profiles of cell biomass (OD_700_), glucose consumption, and CoQ_10_ production during 250 mL shake-flask fermentation of HY12. **f-g,** Fed-batch fermentation of HY12 in a 50-L pilot-scale bioreactor. Time-course monitoring of CoQ_10_ production, dry cell weight (DCW), residual glucose, and pH. Pilot-scale fermentation was carried out, and data were collected at Inner Mongolia Kingdomway Pharmaceutical Co., Ltd.

HY12 (double deletion of RSP_1028&RSP_3509 in HY11) displayed the largest gains, with shake-flask titers matching 48-well plate results. Profiling indicated slower initial HY12 growth (OD_700_) vs. HY01 (0-36 h; *P* < 0.05) but greater biomass later (36-72 h; Fig. 5c). Glucose uptake followed suit, with early reduction (P < 0.05; Fig. 5d). CoQ_10_ titers exceeded HY01 from 36 h, by 36-39% at 60-72 h (P < 0.05; Fig. 5e), denoting enhanced robustness. In 50 L bioreactor scale-up (controlled DO, pH, fed-batch), HY12 reached 3.6 g/L CoQ_10_ at 90 h, 32% above HY01. This titer ranks among the highest reported for microbial CoQ10 production, confirming the strategy for industrial application.

## Discussion

This study unveils physiological mechanisms in the regulatory network governing CoQ_10_ in the photosynthetic bacterium *R. sphaeroides*, while developing strategies to optimize yield and stability in the HIS *R. sphaeroides* HY01. By integrating omics analyses, transcription factor engineering, and genome-scale CRISPRi, we increased CoQ_10_ production by 38% and stabilized the high-yield phenotype, achieving a titer of 3.6 g/L in 50-L fed-batch fermentation. This framework provides a paradigm for sustainable microbial manufacturing.

We demonstrate that the PrrAB two-component system serves as a core regulator of CoQ_10_ biosynthesis, tightly linking it to photosynthetic physiology. Transcriptomics of the spontaneous mutant HY02 showed downregulation of photosynthetic genes (e.g., bacteriochlorophyll and terpenoid pathways) and upregulation of the respiratory chain (Fig. 1b-c), concomitant with loss of photosynthetic function (Fig. 1h-I). PrrA knockout in HY01 recapitulated the white phenotype and an 84% drop in CoQ_10_ (Fig. 1f), with complementation completely restoring accumulation (Fig. 1g). Studies in wild-type strain 2.4.1 confirm that PrrA regulates approximately 25% of the transcriptome, including direct control of genes involved in electron transport, tetrapyrrole synthesis, and terpenoid backbone biosynthesis ^31, 36^. Our work extends PrrAB to industrial contexts, revealing its control of CoQ_10_ as a quinone electron carrier in shared electron transport chains, maintaining energy allocation during microaerobic fermentation^37^. The strong correlation between bacteriochlorophyll and CoQ_10_ (R² = 0.78; Fig. 2a) suggests PrrA-mediated flux partitioning for adaptation to low-oxygen or anoxic stress ^38^. This correlation also supported phenotype-based screening, bypassing detection bottlenecks for membrane-localized CoQ_10_.

By employing the gTME strategy, this study introduces random point mutations in PrrA to enable further CoQ_10_ accumulation. Overexpression of PrrA with strong promoters (*Prsp_7571* or *tac*) ^25^ in HY01 failed to increase CoQ10 production (Fig. 1g), underscoring the need for multi-pathway rebalancing. We previously constructed a *PrrA* mutant library and screened for CoQ_10_-overproducing strains based on greener phenotypes. However, visual differentiation from the parental strain HY01 was low-efficiency, and chemical quantification of bacteriochlorophyll is labor-intensive ^39^ and unsuitable for high-throughput screening. An error-prone PCR library combined with an RFP reporter system (driven by bchC/E promoters ^31^; Fig. 2b-d) identified activating mutations E76G and D116K, each boosting ∼15% (Fig. 2e-f). AlphaFold3 modeling localized these sites at the PrrA-PrrB interface, with E76 modulating electrostatics and D116 reshaping hydrogen bonds (Fig. 3a-b). Saturation mutagenesis revealed limited beneficial variants, but synergistic combinations (e.g., E76F&D116T) elevated >30% in deep-well plates (Fig. 3c-d). These modifications optimize phosphotransfer, enhancing PrrA activation under microaerobic conditions to sustain redox balance ^40^. The strategy suits CoQ_10_ "black-box" pathways, where precursor flux and electron dynamics elude rational design ^41^.

Phenotypic instability in the engineered strain HY11 manifested as white colony reversion during liquid passaging (Fig. 3e), arising from CoQ_10_ overaccumulation-induced stress that promotes mutations—a common issue in HISs^9^. CRISPRi screening (Fig. 4a) enriched 10 target genes, involving DNA replication/repair (*e.g.*, RSP_1028, RSP_0524) and 5 genes of unknown function (*e.g.*, WGR-domain RSP_1085; Fig. 4b-c). These results indicate that CoQ_10_ overaccumulation imposes a metabolic burden, probably triggering genomic instability via DNA damage or replication stress, as observed in burdened bacterial ^6^. The WGR-domain protein (RSP_1085) may participate in ADP-ribosylation-like responses to DNA breaks or oxidative stress, similar to PARP-related mechanisms in bacteria ^42^. Genes encoding hypothetical proteins may suggest some unknown adaptation pathways, requiring further functional dissection to elucidate their roles in stress resilience ^43^. Knockouts validated maintenance of the green phenotype and high yield (Fig. 4d-e). Unknown genes may alleviate oxidative damage or genome remodeling under stress. Combinatorial knockouts (e.g., RSP_1028&RSP_1085) stabilized and amplified yields (Fig. 5a-b), implying synergy in burden relief, such as suppressing error-prone repair.

HY12 exhibited adaptive physiological dynamics during fermentation, with limited early growth followed by a late anaerobic surge in biomass and CoQ_10_ production (Fig. 5c–e), consistent with PrrAB-mediated oxygen sensing that prioritizes anaerobic metabolic flux ^44, 45^. Pilot-scale fermentation achieved 3.6 g/L CoQ10 (Fig. 5f), surpassing previously reported yields of 1–3 g/L ^17^. This framework is applicable to other stress response pathways, offering significant potential for industrial applications. Future multi-omics analysis of novel genes will elucidate adaptation mechanisms in photosynthetic bacteria, advancing sustainable biotechnology.

In summary, we devised a multifaceted engineering approach that integrates comparative omics, structure-guided protein engineering, and high-throughput genetic screening to yield a stable *R. sphaeroides* strain with markedly enhanced CoQ_10_ production, reaching record titers in pilot-scale fermentations. Our results illustrate how precise modulation of oxygen-sensing regulators and targeted disruption of instability-associated pathways can reconcile metabolic productivity with enduring strain fitness, establishing a versatile framework for engineering resilient photosynthetic microbes toward the sustainable manufacture of high-value compounds.

## Methods

### Strains, plasmids, and culture conditions

The strains and plasmids used in this study are Listed in Supplementary Tab.1 and Tab.2. *E. coli* DH10b, *E. coli* S17-1 and their derivatives were cultivated in Luria–Bertani medium at 37°C for plasmid construction and di- parental conjugation. *R. sphaeroides* and its derivatives were cultivated on agar plates (0.8% yeast extract, 0.2% sodium chloride, 0.13% monobasic potassium phosphate, 0.0125% magnesium sulfate, 0.3% glucose and 2% agar, supplemented with 1 mg/L nicotinic acid, 15 mg/L biotin and 1 mg/L thiamine hydrochloride). For 48-well plate, shake-flask or 50-L bioreactor fermention, *R. sphaeroides* and its derivatives were cultivated in fermentation medium (0.4% corn steep liquor, 0.28% NaCl, 0.63% MgSO4, 0.3% sodium glutamate, 0.3% KH_2_PO_4_, 0.3% (NH_4_)_2_SO_4_, 4% glucose and 0.2% CaCO3 supplemented with 1 mg/L thiamine hydrochloride, 1 mg/L nicotinic acid, and 15μg/L biotin).

### RNA sequencing (RNA-seq) and transcriptome analysis

RNA-seq was performed as described previously ^3^. Total RNA was isolated using a Redzol reagent kit from SBS Genetech Co. Ltd (Beijing, China). The quality of the RNA samples was analyzed using an Agilent Bioanalyzer 2100 system (Agilent Technologies), and mRNA was enriched by rRNA depletion and followed by mRNA fragmentation, cDNA strand synthesis and library construction. The RNA-seq and transcriptomic analyses were performed by Novogene Co., Ltd (Beijing, China).

### Structural modeling

PrrA is the response regulator (RR) of the two-component system PrrAB, binding to target DNA sites and activating gene transcription when phosphorylated at aspartate 63. To further understand how PrrAE76G affects its regulation property, structural prediction of protein-protein interaction (PPI) was explored on AlphaFold 3 sever ^46^. We hypothesized that if the E76G could alter the PrrAB interaction and thereby impact downstream signaling. To test this, we submitted 5 parallel jobs to AlphaFold 3 sever with random seeds using the sequences of two copies of PrrB and two copies of either PrrAWT or PrrA mutant. Fifty molecules of oleic acid were also included in each prediction to mimic the membrane environment, as the N-terminus of PrrB contain a transmembrane region.

### Genetic manipulation and gene overexpression in *R. sphaeroides*

The primers used for plasmids construction are present in Supplementary Tab. 3. For *mppG*, *ppsR*, *appA*, *tspO*, *crpK*, *prrA*, *rsp_0524*, *rsp_1028*, *rsp_2679*, *rsp_1085*, *rsp_1864*, *rsp_1781*, *rsp_2382*, *rsp_2675*, *rsp_2075* and *rsp_3509* gene deletion in *R. sphaeroides*, plasmid pK18mobsacB (pK18mob) was digested by HindIII and BamHI to obtain a linear vector, and a high fidelity KOD one polymerase (Bio-toyobo, Inc.) was used to amplify two ∼500 bp homology arms by using *R. sphaeroides* genomic DNA as template. The linear vector DNA and PCR products were assembled with a Seamless Assembly Mix Kit (Abclonal Technology, Inc.). A pBBR1MCS-NEW derivative was used for gene overexpression. Targeted genes *prrA* were amplified from *R. sphaeroides* HY01 genomic DNA. Conjugation was performed as described previously ^25^. The plasmid was introduced into *R. sphaeroides* with the help of *E. coli* S17-1 by intergeneric conjugation, and exconjugants with single-crossover were selected for kanamycin resistant (kan^R^). Then, the mutant was selected from the initial exconjugants after several rounds of nonselective growth, and confirmed by PCR amplification using the primers list in Supplementary Tab. 4.

### Fermentation in Shake-flasks

The *R. sphaeroides* strains were cultivated in 250 mL shake-flasks with 25 mL of seed medium at 32 °C and 220 rpm for 16-18 h. Subsequently, the seed culture was inoculated into 250-mL shake flasks containing 45 mL of fermentation medium with a starting optical density (OD_700_) of 1.2. The pH was adjusted to 7.0 with a NaOH solution. *R. sphaeroides* cells were grown in a fermentation medium at 32 °C with 220 rpm for 72 h, and 1 mL of sample was collected every 12 h.

### High-Performance Liquid Chromatography (HPLC) detection of CoQ_10_

CoQ_10_ was measured by HPLC. First, 1 mL of fermentation broth was mixed with 10 μL of 6 N HCl and 0.2 mL 30% hydrogen peroxide, followed by the addition of 2 mL acetone and vertexing for 1 min. The volume was subsequently adjusted to 10 mL with ethanol, followed by incubation in an ultrasonic bath for 45 min at room temperature, and centrifuged at 12,000 rpm for 10 min (#37520, Thermo Fisher Scientific, Inc.). Then the supernatant was filtered using a 0.45-μm membrane (Merck Millipore). The resulting samples were then used for CoQ10 detection by HPLC. A YMC-Pack ODS-A C18 column (150 mm × 4.6 mm; YMC Co., Ltd., Tokyo, Japan) for HPLC analysis on an Agilent 1260 system (Agilent Technologies, Santa Clara, CA, USA). The mobile phase (methanol: ethanol; 65: 35) was applied at a flow rate of 1.5 mL/min at room temperature, and the eluate was monitored at 275 nm using a photodiode array detector (Agilent Technologies, Santa Clara, CA, USA).

### CRISPR/dCas12a-mediated CRISPRi and genome-wide transformants screening

To identify novel targets associated with strain stability, 100-200 μL of HY11 strain containing the CRISPRi library was inoculated into 25 mL seed medium (25 mg/mL kanamycin) and incubated at 32°C with 220 rpm agitation for 18-24 hours until the OD_700_ reached approximately 8-10. A 10 mL aliquot (20%) of the seed culture was then inoculated into a fermentation medium (25 mg/mL kanamycin and 0.2-0.4 mM IPTG) and subjected to shaking culture at 32°C with 220 rpm for 48 hours. Upon completion of fermentation, 1 mL of the culture was serially diluted to 10^-1^, 10^-2^, …, 10^-7^, and 100-200 μL aliquots of each dilution were spread onto plates containing 25 mg/mL kanamycin and 0.3 mM IPTG. The plates were incubated at 32°C for 5-7 day. Colonies exhibiting a dark green phenotype were selected and subjected to fermentation to assess changes in CoQ_10_ production. PCR amplification and sequencing were subsequently performed to confirm the inhibitory targets.

### Fed-batch fermentation

Fed-batch fermentation of CoQ10 was performed in a 50-L stirred bioreactor (Shanghai Guoqiang Bioengineering Equipment CO., LTD, Shanghai, China) with an initial working volume of 18 L. Foam formation was prevented by the addition of antifoam 204 (Sigma–Aldrich, St. Louis, MO, USA). Dissolved oxygen (DO) and pH were measured by a DO probe (VisiFerm DO Arc 225 H2, Hamilton, Switzerland) and a pH electrode (EasyFerm Plus PHI Arc 225, Hamilton, Switzerland), respectively. The temperature was maintained at a constant 32°C, aeration at 1.0 VVM, agitation at 600 rpm, and pH at 6.5 by automatic injection of acetic acid or ammonia. The fed-batch process was initiated after 16 h of cultivation from a 500 g/L concentrated glucose stock solution.

### Cell growth and sugar analysis

Growth of *R. sphaeroides* cells was detected by measuring the optical density at 700 nm (OD_700_). Initially, 0.5 mL of culture broth was mixed with 0.2 mL of 0.1 N HCl to completely dissolve CaCO_3_, followed by dilution with deionized water and measurement of the OD_700_ using a spectrophotometer. Residual glucose in the culture broth was measured using an SBA-40D biological sensing analyzer (Biology Institute of the Shangdong Academy of Science, Jinan, China) according to manu-facturer instructions.

### Statistical analysis

Data analysis and process were performed using Microsoft Excel 2016, Origin 2025. Unless otherwise indicated, data in figures were expressed as mean ± standard deviation (SD). Where relevant, two-tailed Student’s t-tests were used to demonstrate differences in values. *P* < 0.05 was used as a standard criterion of statistical significance.

## Data availability

The authors declare that the primary data supporting the findings of this study are accessible within the paper and its Supplementary Information files.

## Author contributions

G.-Y.T. and L. Z supervised the project. G.-Y.T. and M.L. designed the experiments. M. L., X. H., J. L., K. W., and Y. H. performed the genetic, molecular biology, and laboratory-scale fermentation experiments. D. L. and G. L. performed AlphaFold modeling and related structural analyses. M.L. conducted high-throughput screening experiments. Z. Z., D. L., and B. C. conducted the fed-batch work in 50-L bioreactor. G.-Y.T. and M. L. wrote the manuscript. G.-Y.T., L. Z., L. B. and W. W. edited the manuscript.

## Competing interests

The authors have declared no competing interest.

## Supplementary Information

### Supplementary Figures

**Supplementary Fig. 1.**
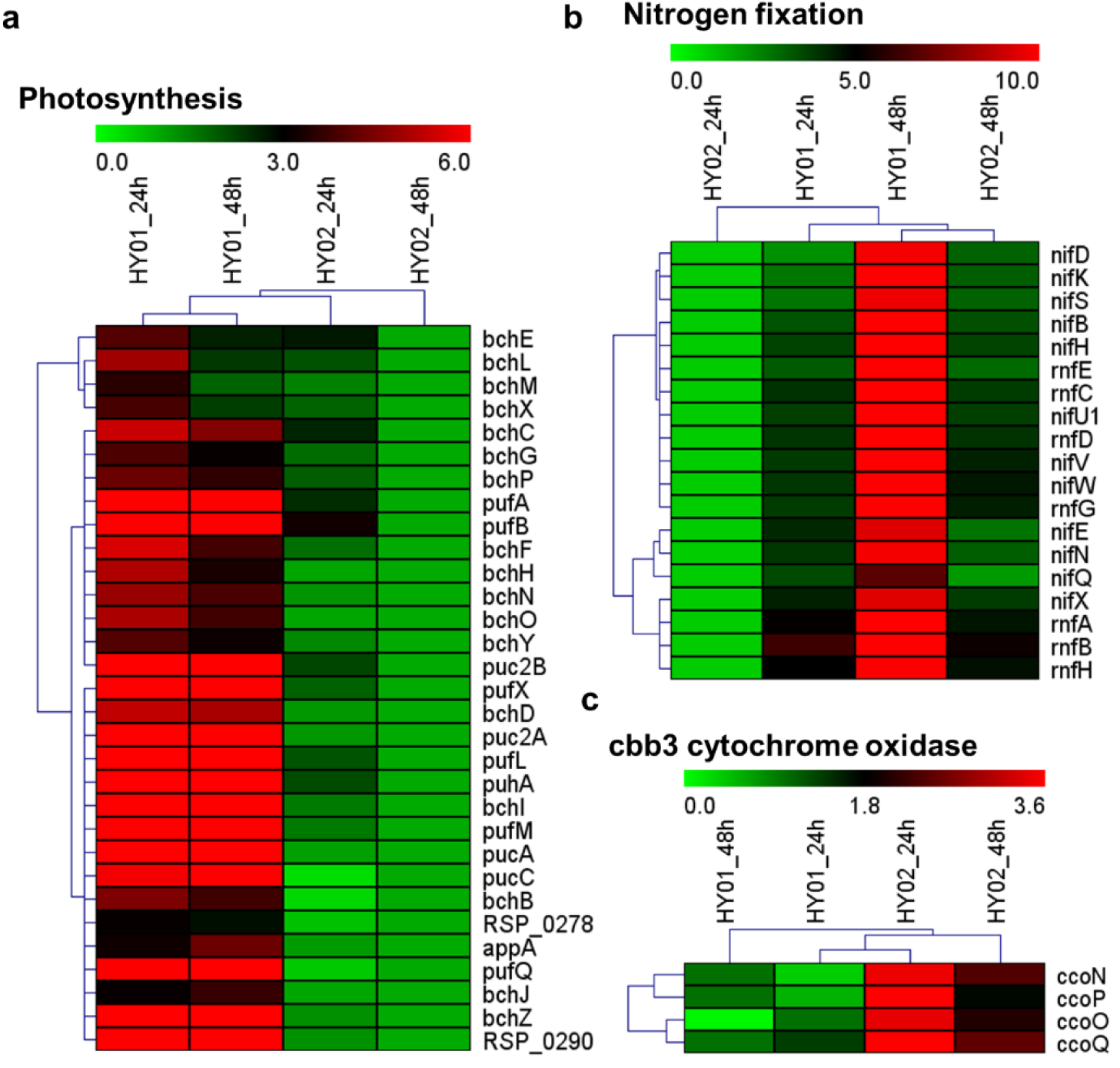
Transcription of genes involved in photosynthesis (a), nitrogen fixation (b), and cbb3- type cytochrome oxidase (c) in HY01 and HY02.

**Supplementary Fig. 2.**
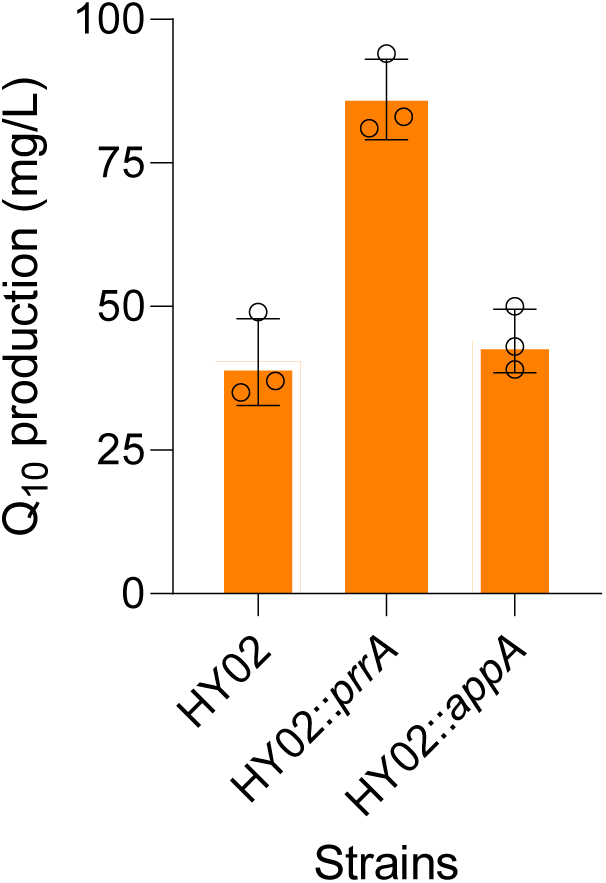
Effects of *prrA* and *appA* duplication on CoQ_10_ production in HY02.

**Supplementary Fig. 3.**
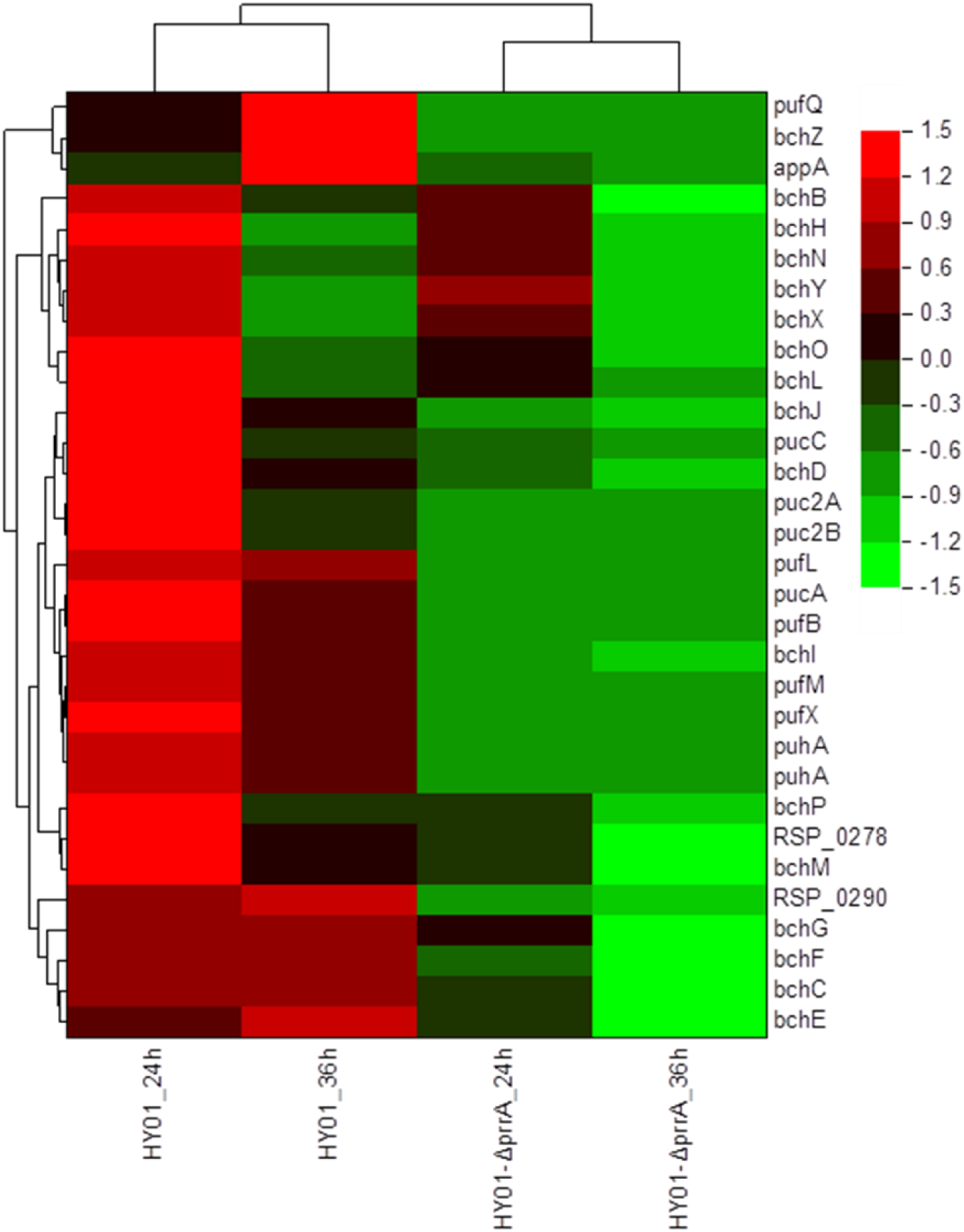
Transcription of genes involved in photosynthesis in HY01-Δ*prrA*.

**Supplementary Fig. 4.**
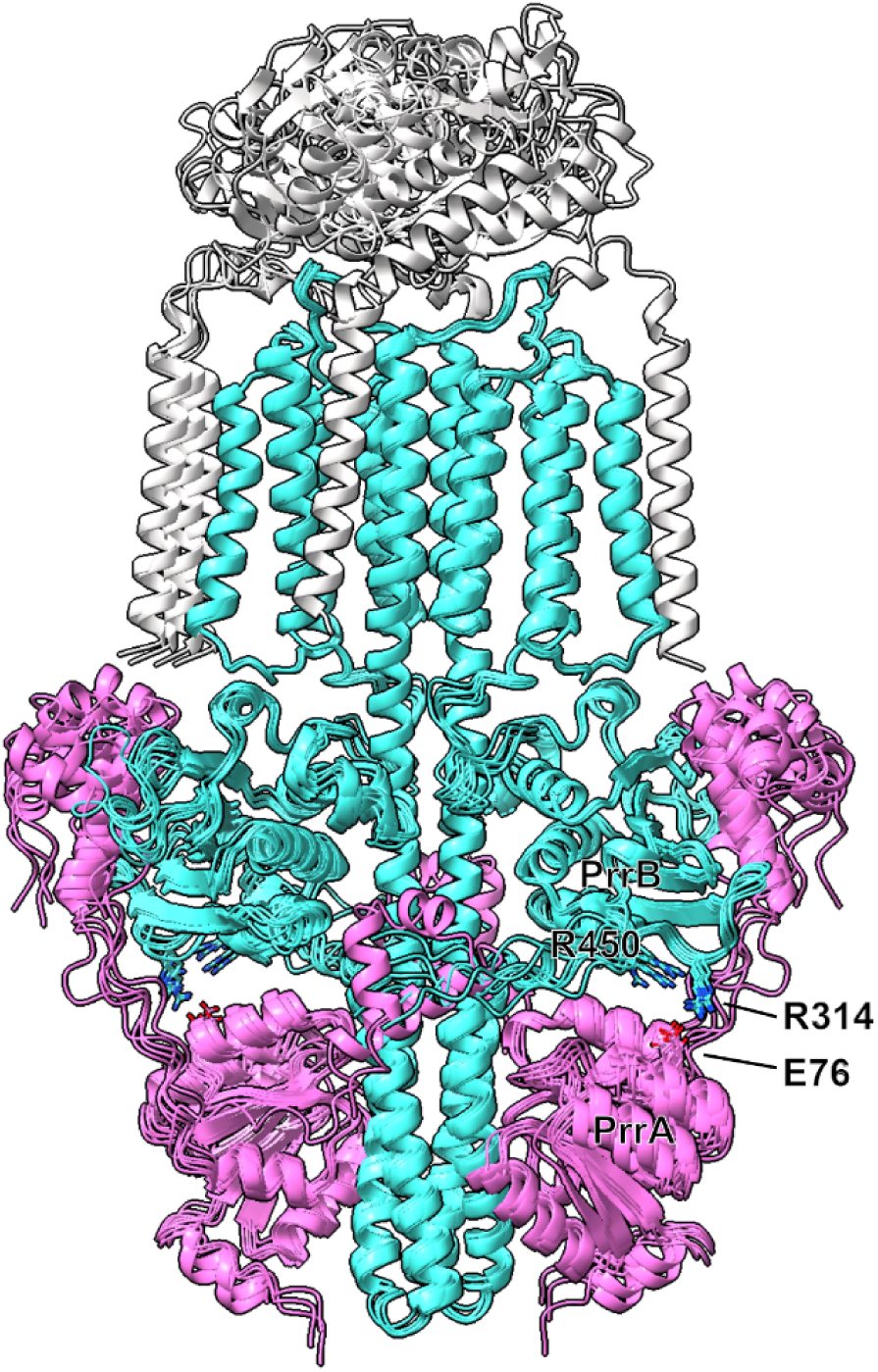
Superposition of five randomly seeded predictions of the 2PrrA–2PrrB–1PrrC complex. PrrA is shown in magenta, PrrB in cyan, and PrrC in grey. Residue E76 of PrrA, and residues R314 and R450 of PrrB are highlighted as ball-and-stick models in their respective colors and labeled.

**Supplementary Fig. 5.**
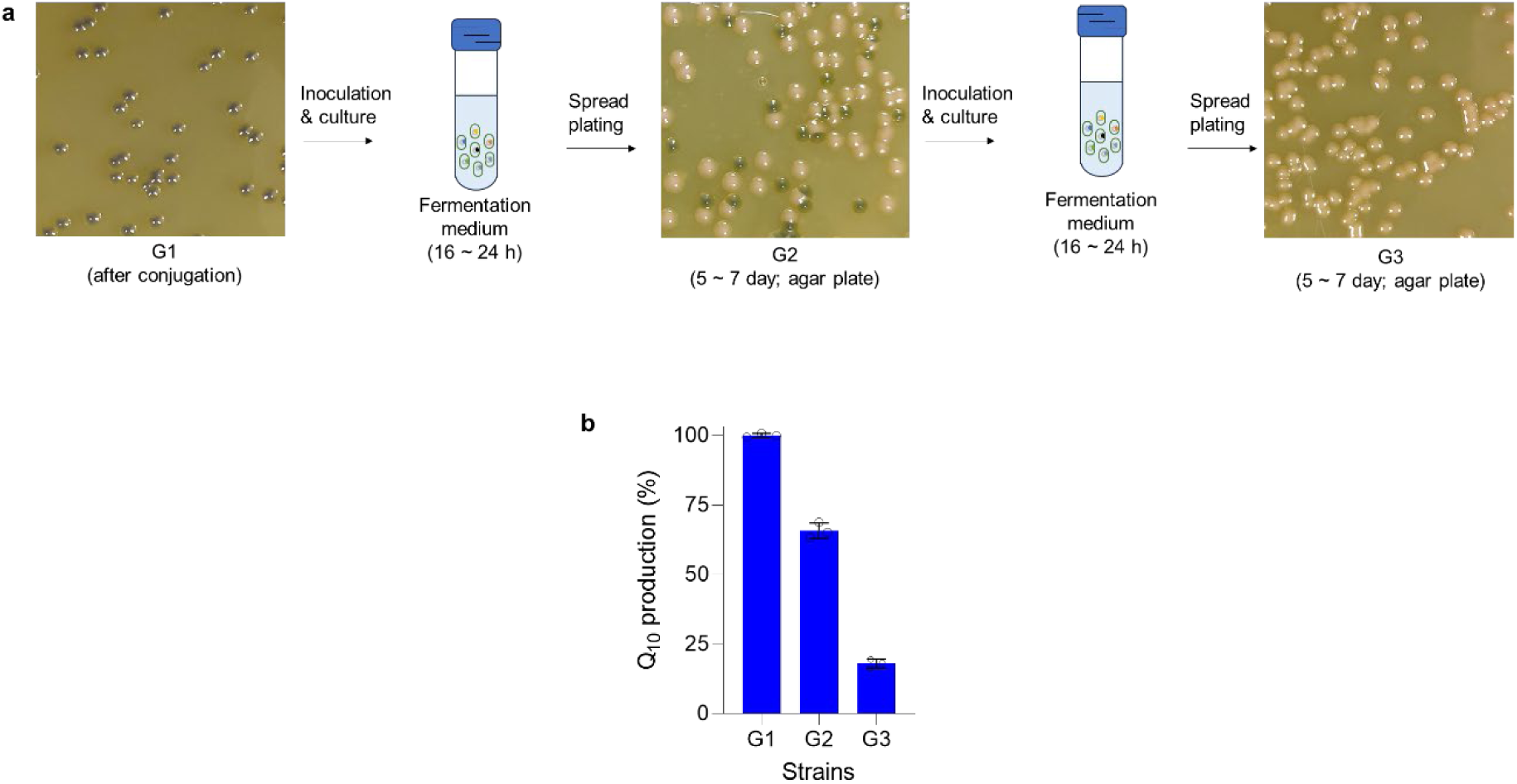
Stability testing of HY11 through continuous liquid fermentation. **a,** Schematic diagram of the continuous liquid fermentation process. **b,** Coenzyme Q10 production at different rounds of fermentation

### Supplementary Tables

**Supplementary Table 1.**
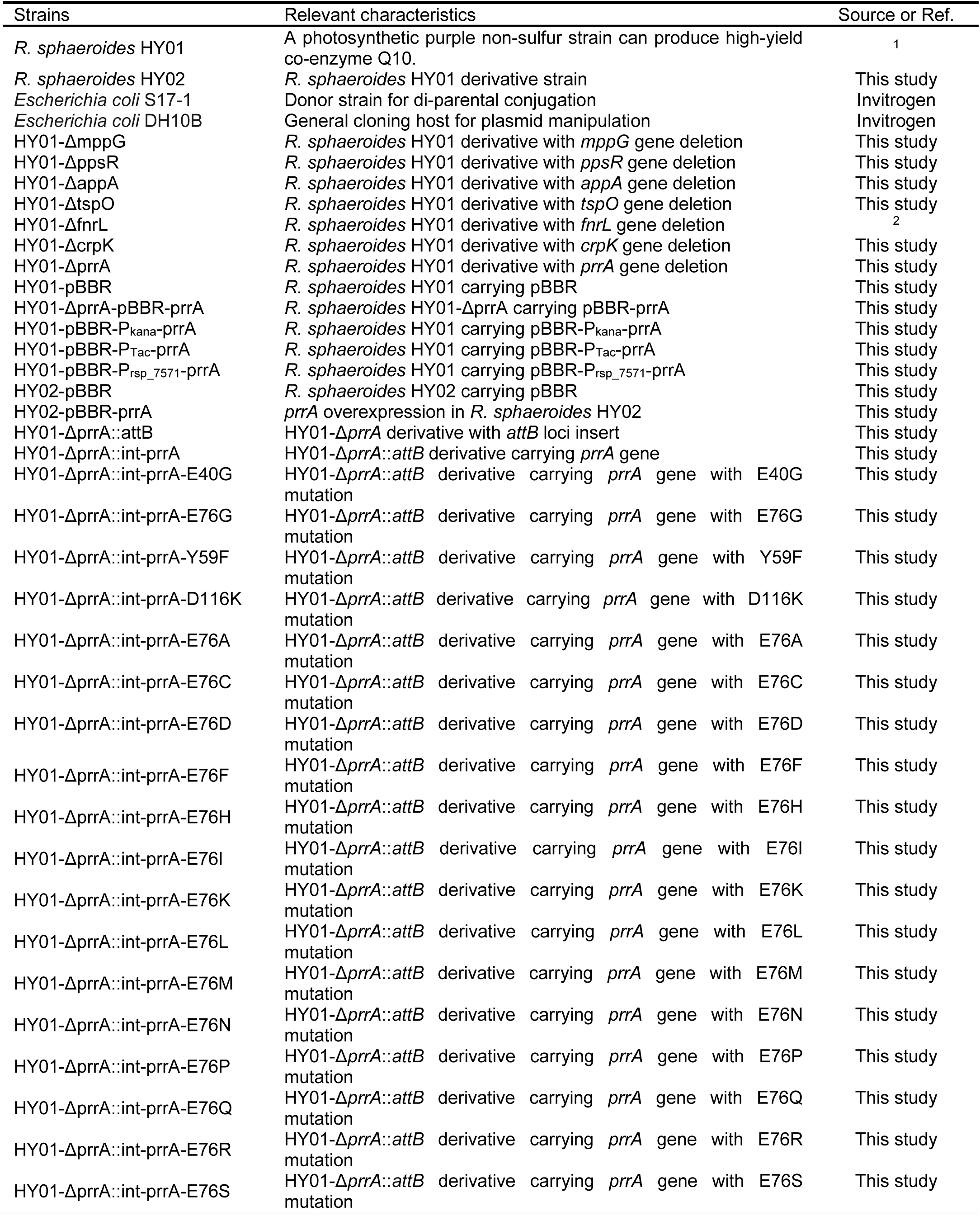
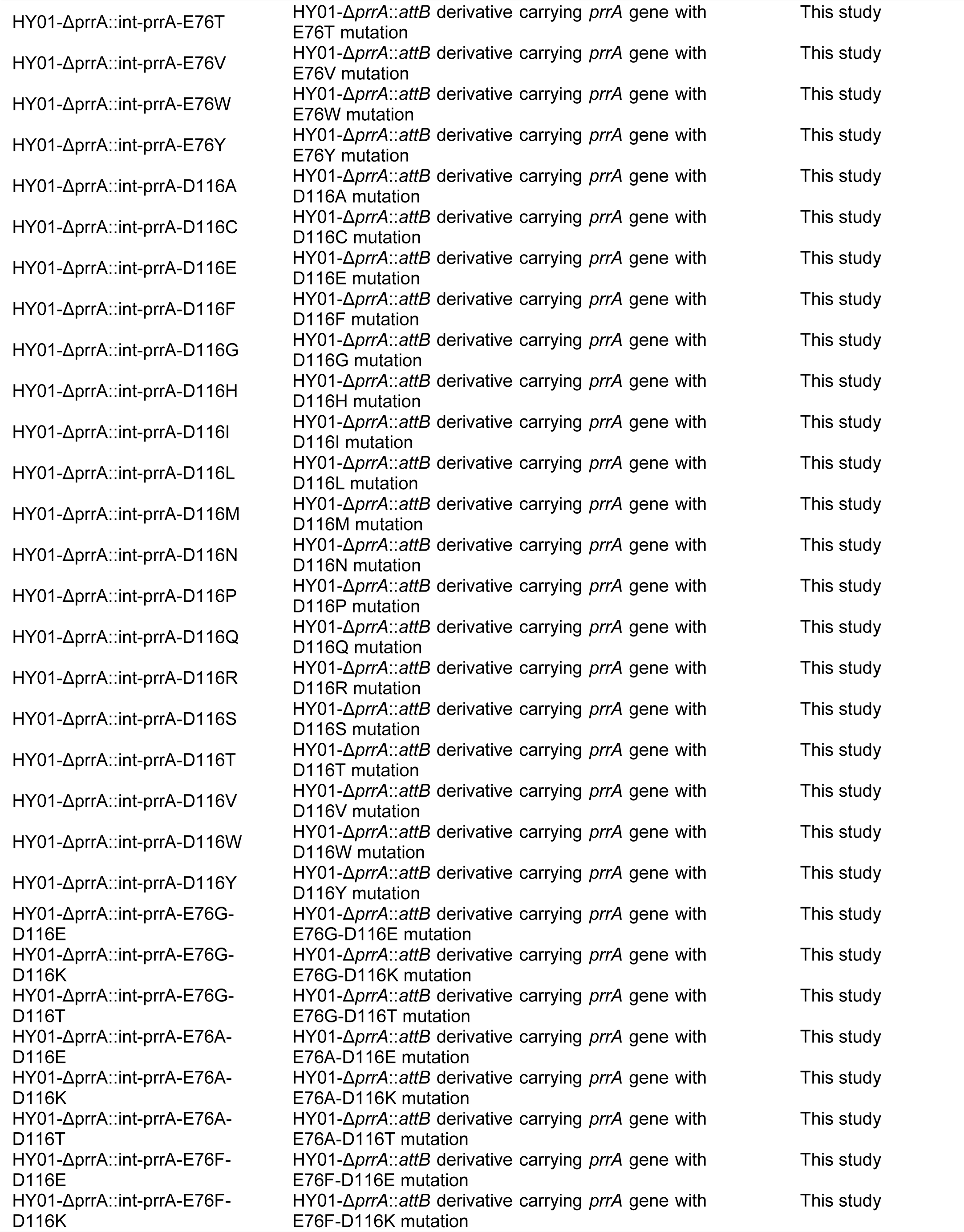
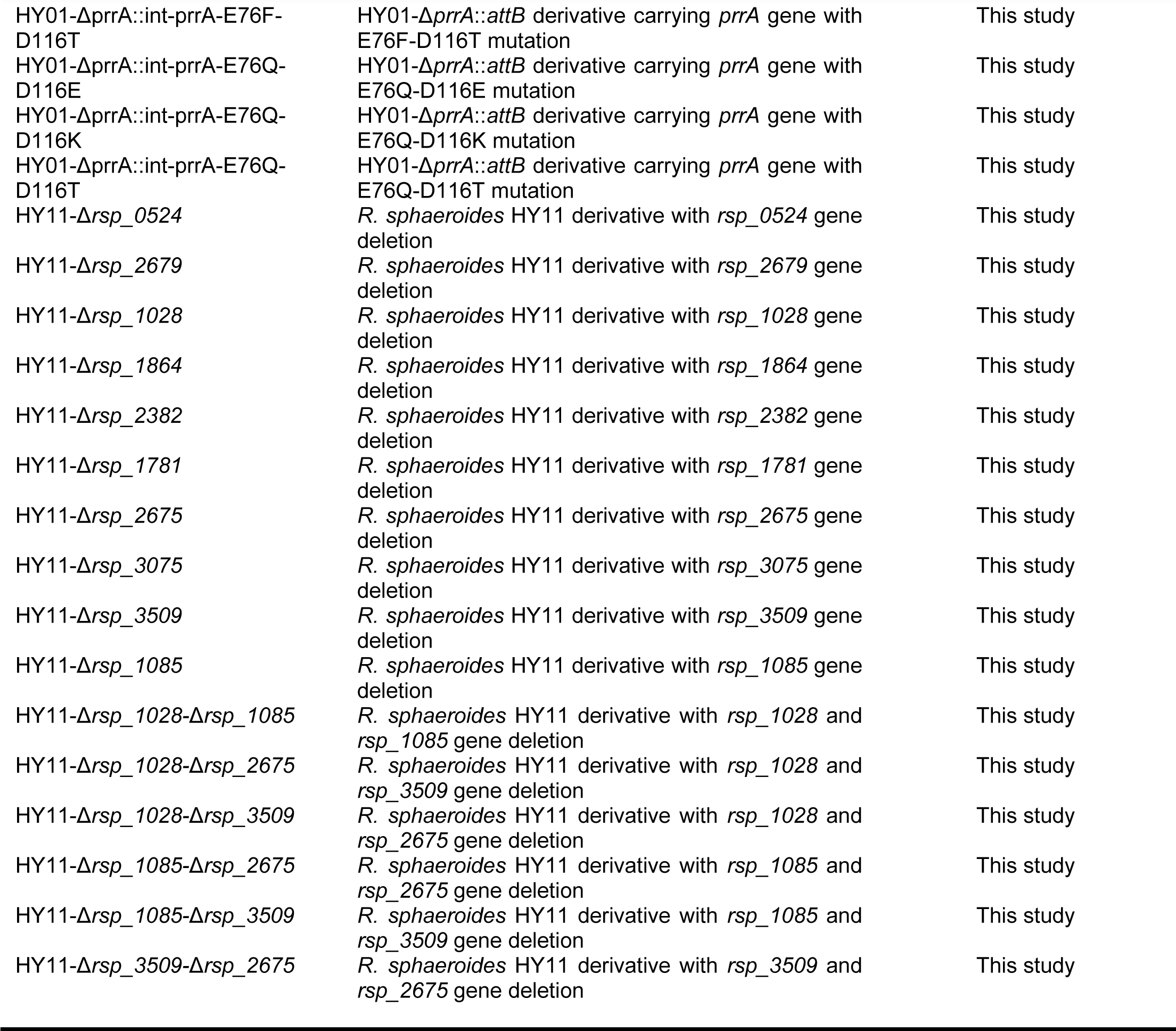
The strains used in this study.

**Supplementary Table 2.**
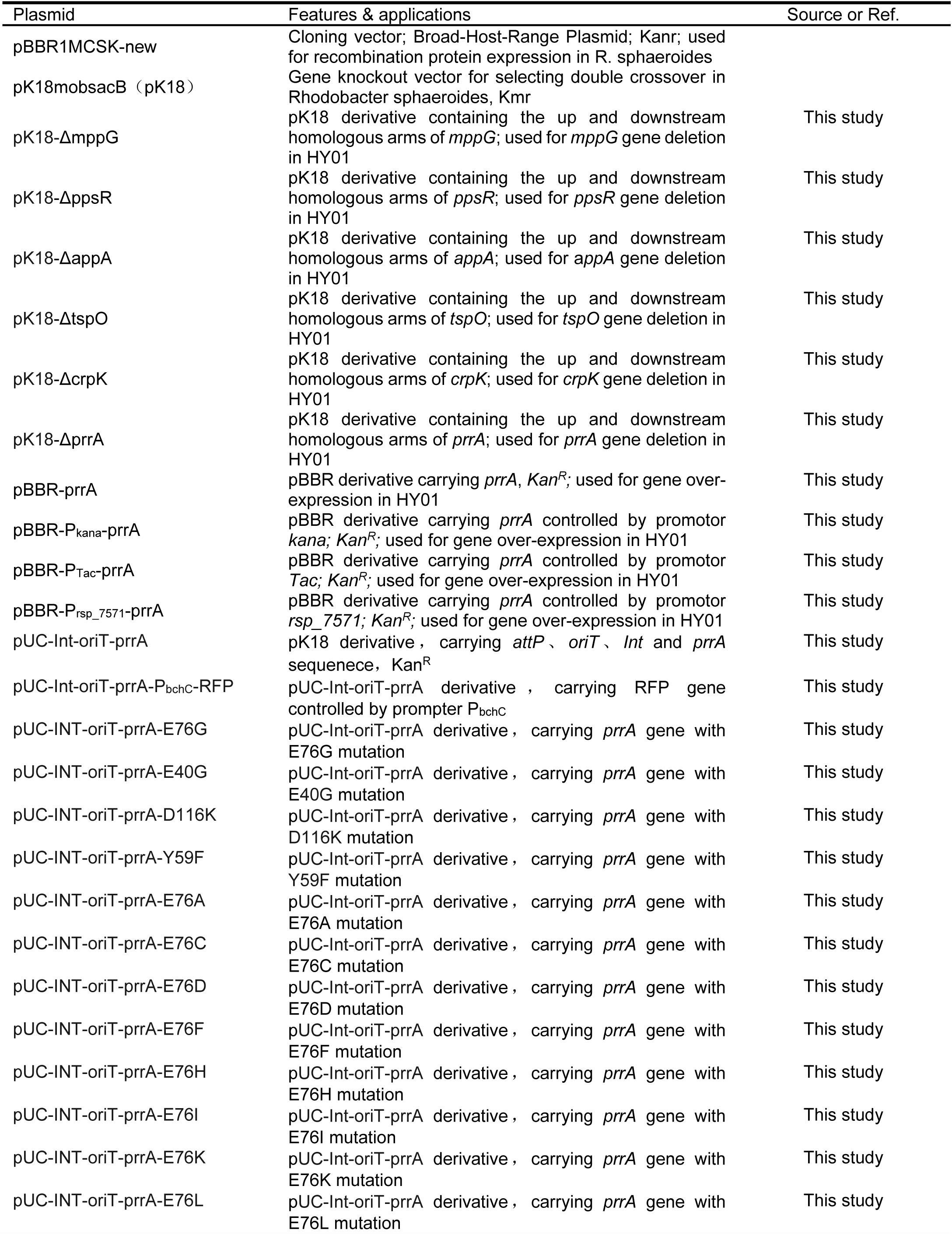
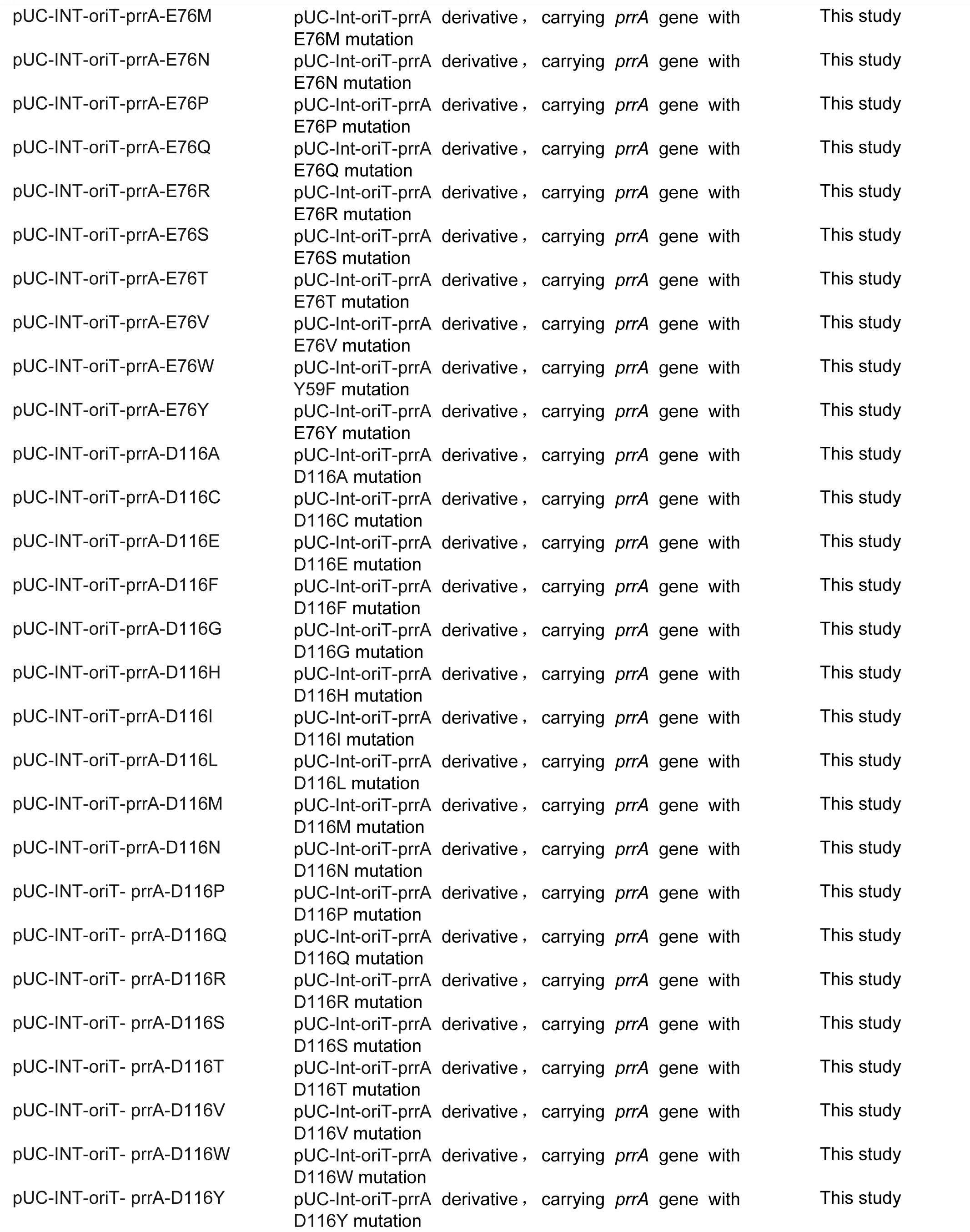
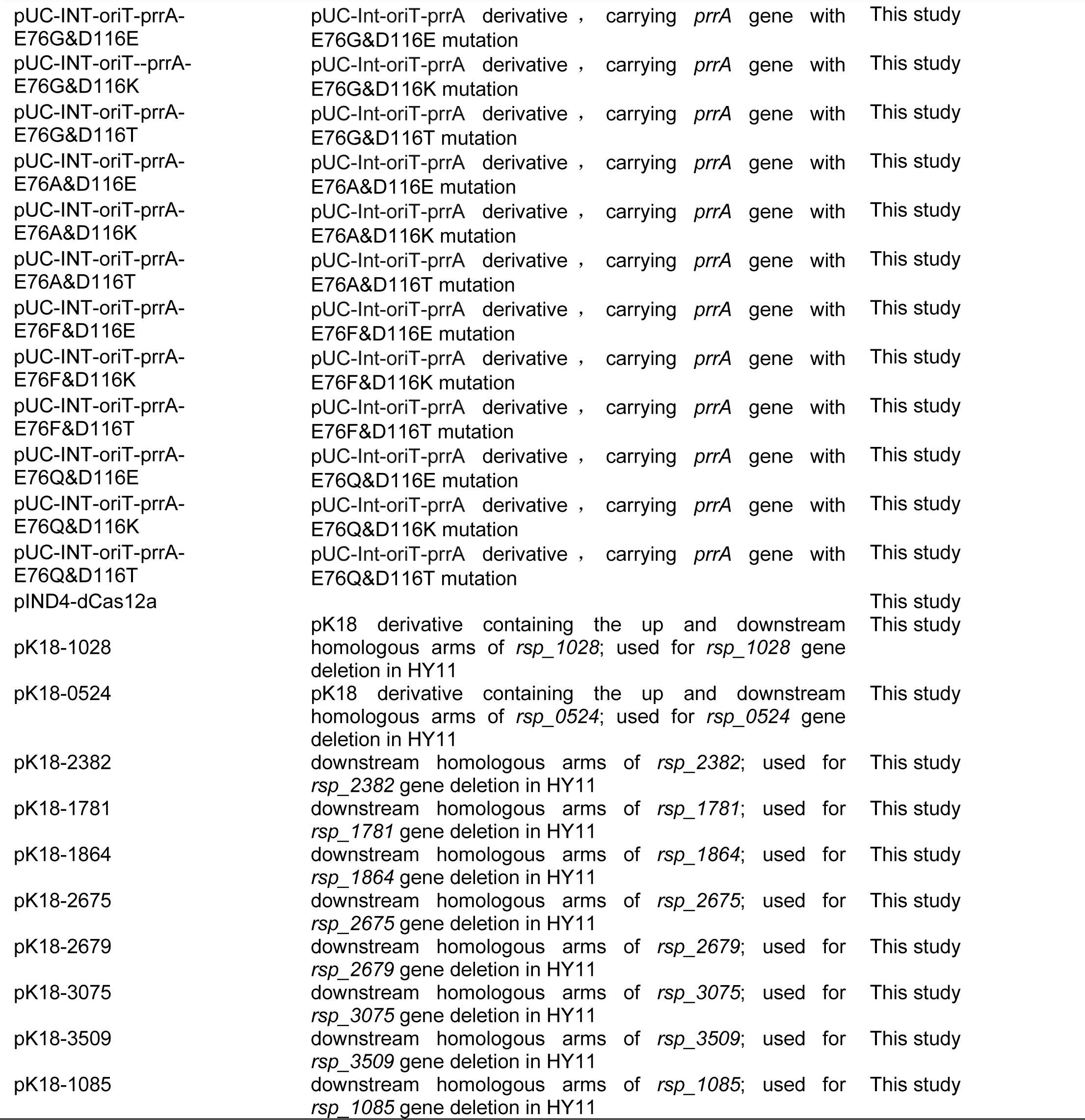
The plasmids used in this study.

**Supplementary Table 3.**
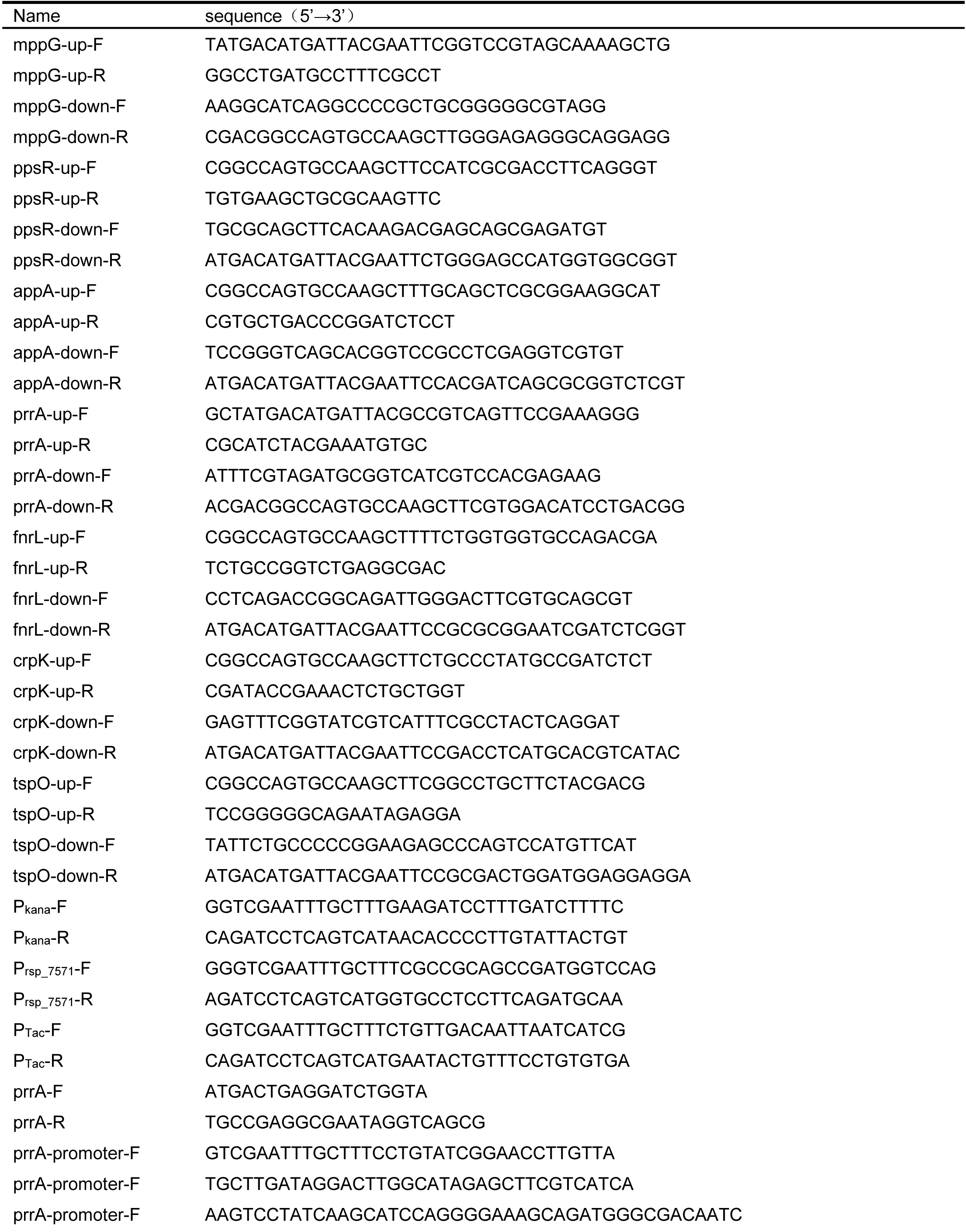
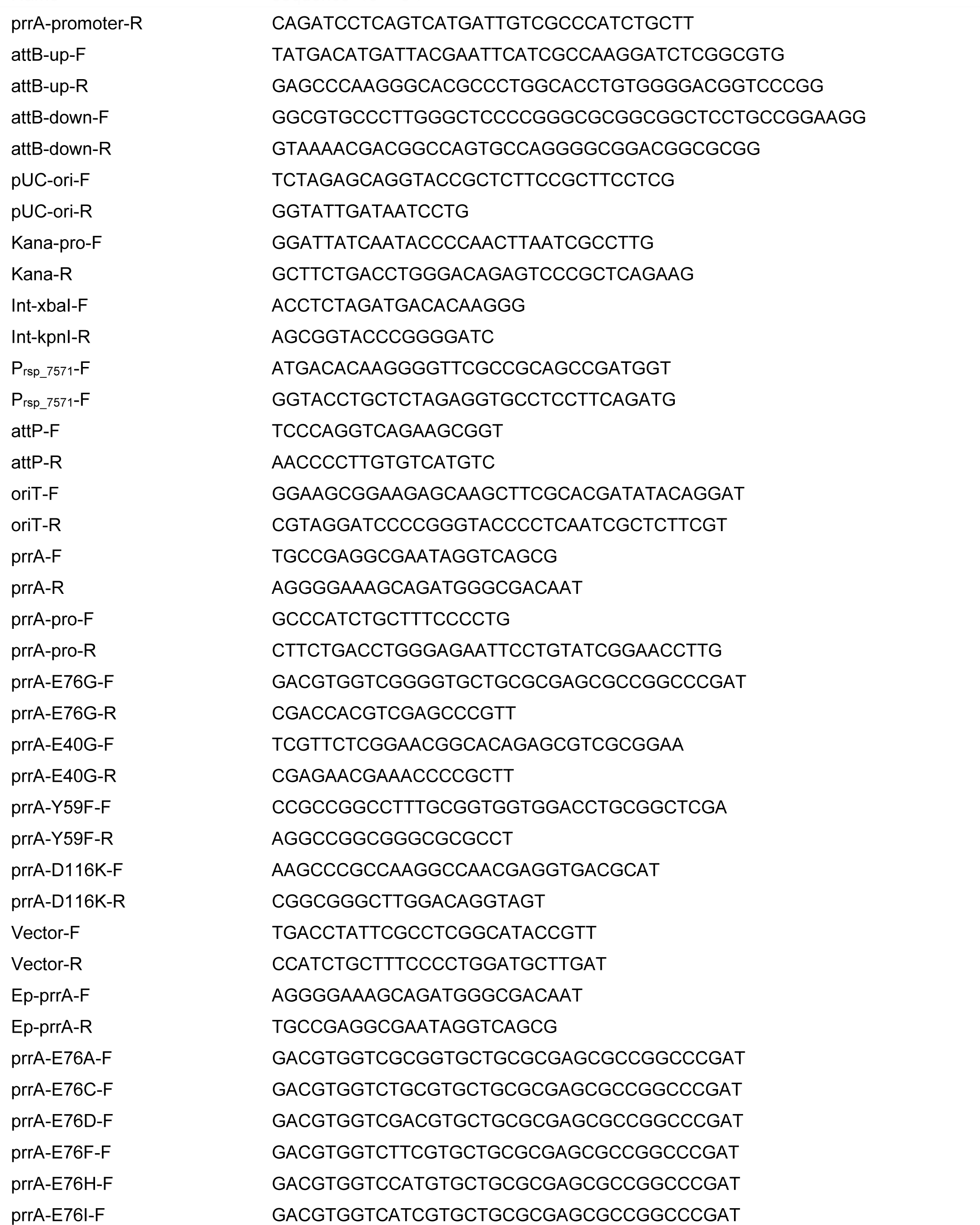
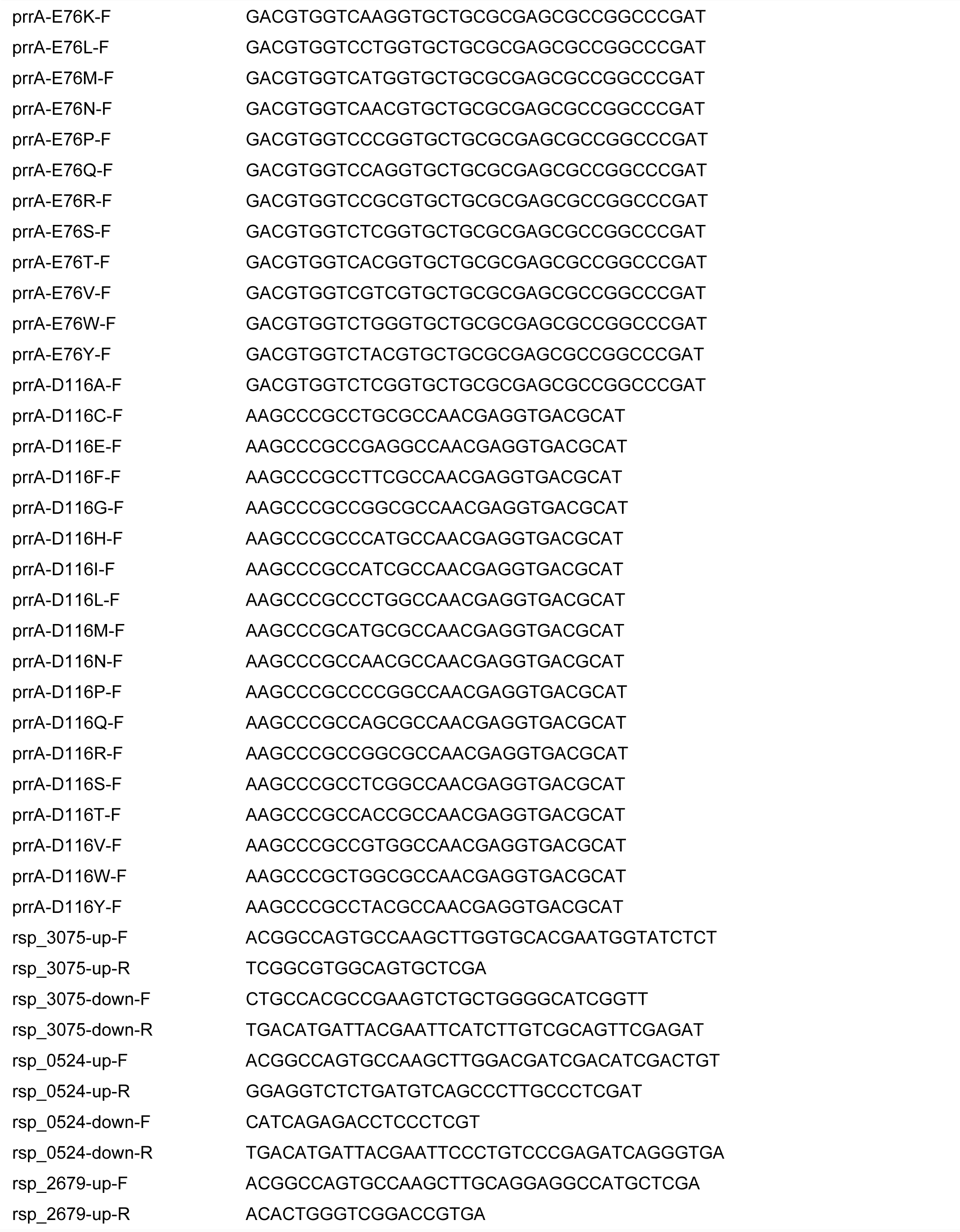
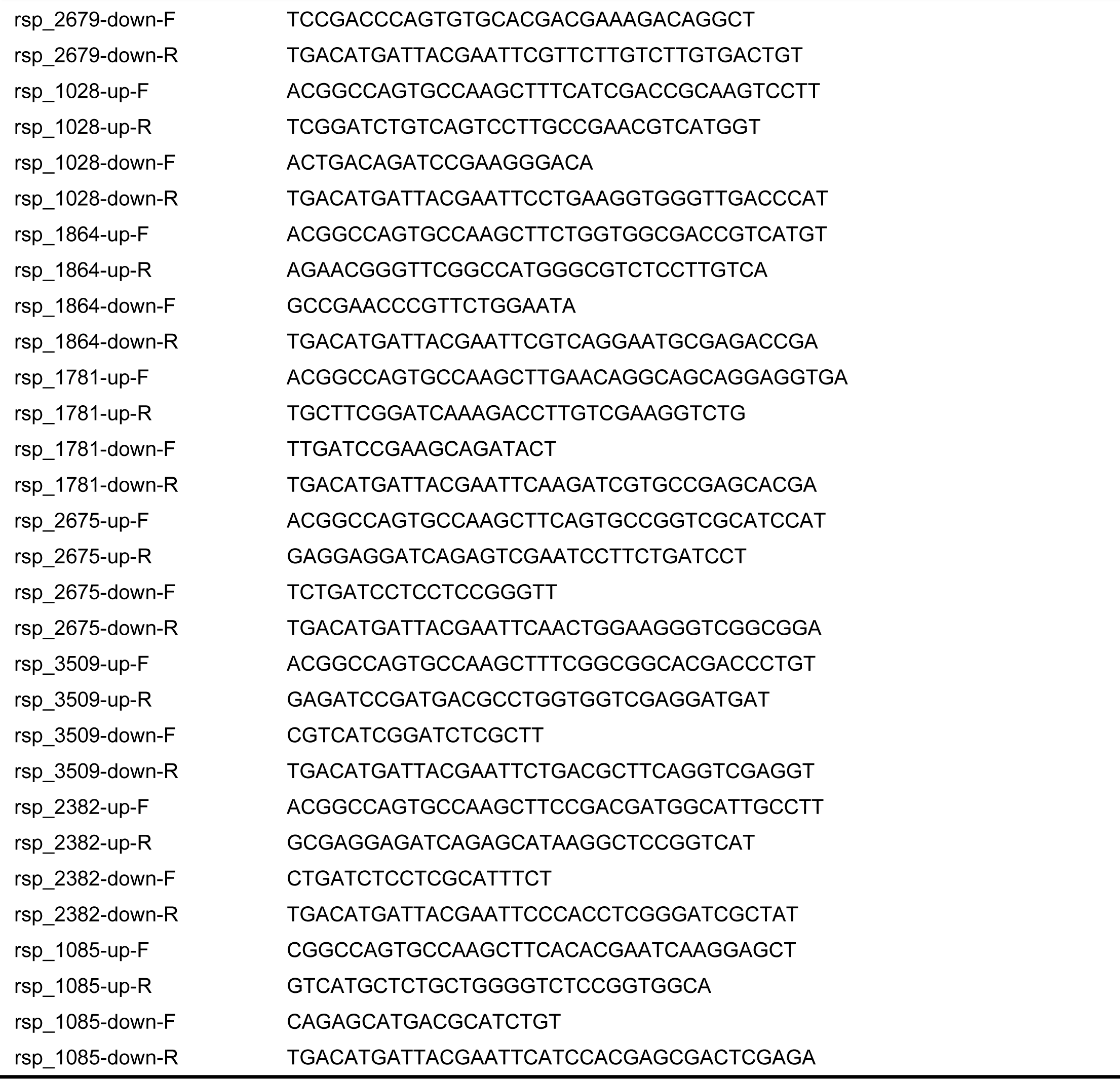
The sequences of primer used in this study.

**Supplementary Table 4.**
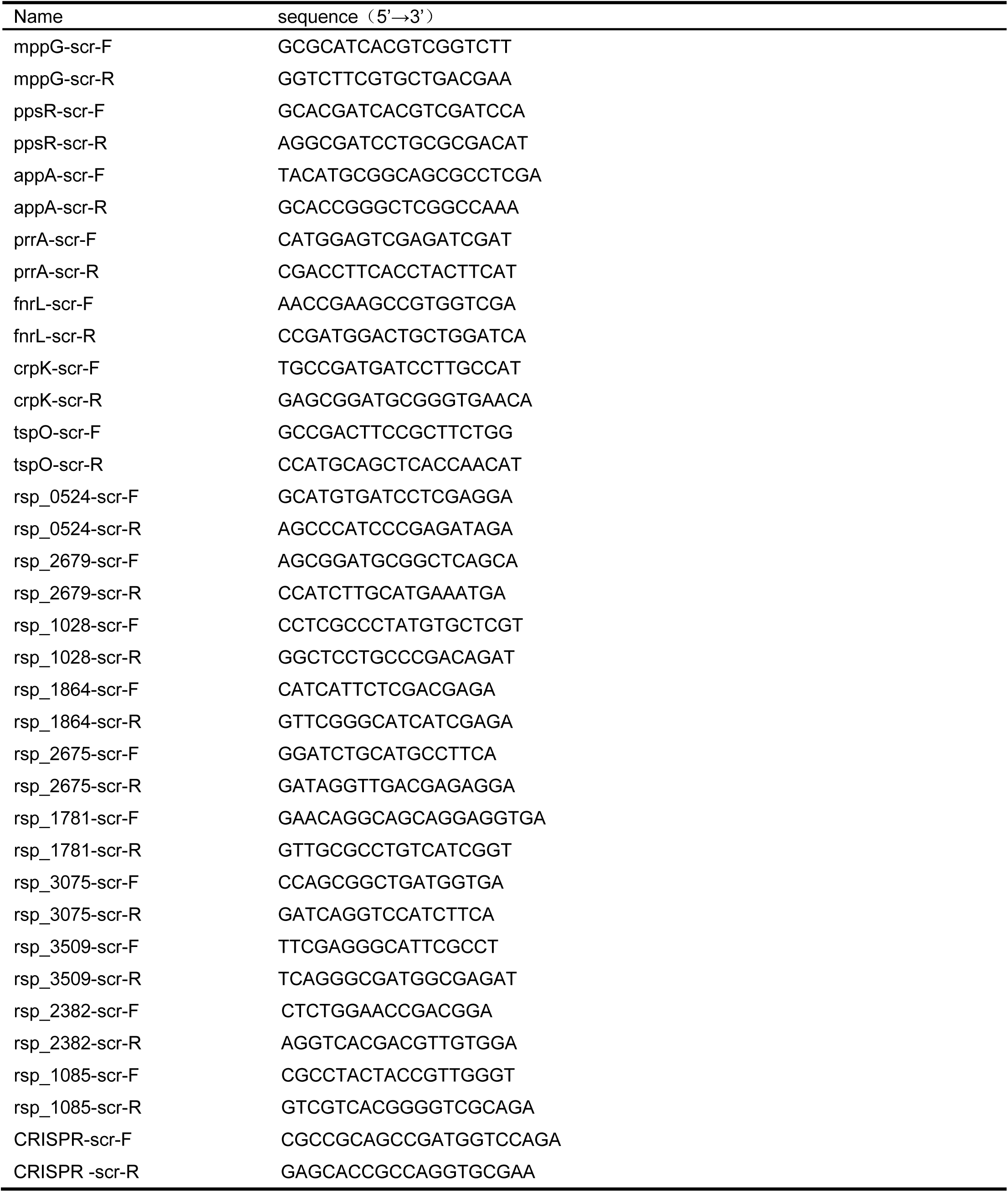
The sequences of screening primer used in this study.

## Notes

### Summary of Updates

The manuscript itself remains unchanged; this resubmission only completes the previously missing author and related information.

